# Signatures of host specialization and a recent transposable element burst in the dynamic one-speed genome of the fungal barley powdery mildew pathogen

**DOI:** 10.1101/246280

**Authors:** Lamprinos Frantzeskakis, Barbara Kracher, Stefan Kusch, Makoto Yoshikawa-Maekawa, Saskia Bauer, Carsten Pedersen, Pietro D Spanu, Takaki Maekawa, Paul Schulze-Lefert, Ralph Panstruga

## Abstract

Powdery mildews are biotrophic pathogenic fungi infecting a number of economically important plants. The grass powdery mildew, *Blumeria graminis,* has become a model organism to study host specialization of obligate biotrophic fungal pathogens. We resolved the large-scale genomic architecture of *B. graminis forma specialis hordei (Bgh)* to explore the potential influence of its genome organization on the co-evolutionary process with its host plant, barley *(Hordeum vulgare).* The near-chromosome level assemblies of the *Bgh* reference isolate DH14 and one of the most diversified isolates, RACE1, enabled a comparative analysis of these haploid genomes, which are highly enriched with transposable elements (TEs). We found largely retained genome synteny and gene repertoires, yet detected copy number variation (CNV) of secretion signal peptide-containing protein-coding genes (*SPs*) and locally disrupted synteny blocks. Genes coding for sequence-related SPs are often locally clustered, but neither the *SP* clusters nor TEs are enriched in specific genomic regions. Extended comparative analysis with different host-specific *B. graminis formae speciales* revealed the existence of a core suite of *SPs,* but also isolate-specific *SP* sets as well as congruence of *SP* CNV and phylogenetic relationship. We further detected evidence for a recent, lineage-specific expansion of TEs in the *Bgh* genome. The characteristics of the *Bgh* genome (largely retained synteny, CNV of *SP* genes, recently proliferated TEs and a lack of compartmentalization) are consistent with a “one-speed” genome that differs in its architecture and (co-)evolutionary pattern from the “two-speed” genomes reported for several other filamentous phytopathogens.

## INTRODUCTION

Powdery mildews (Ascomycota, Erysiphales) are ubiquitous fungal plant pathogens in temperate regions of the world (Glawe 2008). They thrive on the basis of an obligate biotrophic lifestyle, *i.e.,* by retrieving nutrients from living plant cells for fungal growth and reproduction, and have been extensively studied regarding molecular and genetic interactions with both host (Kuhn et al. 2016) and non-host plants (Lipka et al. 2008). Despite advances in the deployment of durable resistance (Kusch and Panstruga 2017), powdery mildews remain a constant threat for economically important crops as they rapidly evade selection pressure resulting from fungicide application (Jones, L. et al. 2014; Tucker et al. op. 2015) and resistance (*R*)-gene mediated immunity (Brown 2015). The barley powdery mildew pathogen, *Blumeria graminis* f.sp. *hordei* (*Bgh*), is a member of the species *Blumeria graminis* that is specialized on its host plant, barley (*Hordeum vulgare*). There are various specialized forms (*formae speciales*) of *B. graminis,* where each *forma specialis* (f.sp.) is capable of infecting the respective host plant species belonging to the grasses (Poaceae) family, including cereals (Wyand and Brown 2003). Within each *forma specialis,* numerous isolates (strains) can be differentiated, primarily based on their respective virulence/avirulence phenotypes on particular genotypes of the host population (Lu et al. 2016). Meanwhile, *B. graminis* has become a model organism to study host specialization of obligate biotrophic fungal pathogens.

With the dawn of next-generation sequencing, several studies provided initial insights in the haploid genomes of powdery mildews and the molecular basis of their obligate biotrophic lifestyle. Indeed, the first genomic studies, coupled with other “omics” approaches (Bindschedler et al. 2016), showed that powdery mildews have experienced the loss of several, otherwise widely conserved Ascomycete genes with functions related to carbohydrate degradation, primary and secondary metabolism (Spanu et al. 2010; Wicker et al. 2013), which could explain their strict dependence on live plant tissue. On the other hand, these genomes harbor an abundance of candidate secreted effector protein (CSEP)-coding genes, which were deemed to be crucial for successful pathogenesis (Pedersen et al. 2012; Wicker et al. 2013). Isolate-specific variants of these powdery mildew CSEPs are recognized by matching intracellular immune receptors, encoded by barley or wheat *R* genes, which are present only in particular genotypes of these cereal hosts (Bourras et al. 2016; Lu et al. 2016; Praz et al. 2017). This demonstrates that at least these CSEPs are targets of the plant immune system and indicates co-evolutionary dynamics underlying interactions between the pathogen and cereal hosts at the population level. Genome sequencing of members of the cereal powdery mildew pathogen, *B. graminis*, led to the understanding that host specialization can occur by hybridization between two reproductively isolated *formae speciales* that multiply on different host species (Menardo et al. 2016) and, possibly, also by “host tracking” or co-speciation (Menardo, Wicker et al. 2017; Troch et al. 2012). Comparative sequence analysis of multiple isolates of both barley and wheat powdery mildew pathogens, *B. graminis* f.sp. *hordei* (*Bgh*) and *B. graminis* f.sp. *tritici* (*Bgt*), revealed that at least their genomes are characterized by an ancient haplotype mosaic composed of isolate-specific DNA blocks, suggesting exceptionally rare outbreeding and dominant clonal reproduction of the haploid fungus in nature (Hacquard et al. 2013; Wicker et al. 2013).

Powdery mildew fungi have some of the largest genomes among plant-pathogenic Ascomycetes, strongly enriched with an unusually high content of transposable elements (TE) (Spanu et al. 2010; Wicker et al. 2013). Extensive repetitive regions made up of TEs have hindered high quality short read-based genome assemblies, resulting in severely fragmented datasets (Hacquard et al. 2013; Jones, L. et al. 2014; Spanu et al. 2010; Wicker et al. 2013). The fragmentation of the available genomic assemblies has so far hampered our ability to address relevant biological questions, as for example the existence of long lineage-specific virulence regions, the impact of TEs on genome organization and evolution, as well as the conservation of gene order between diverged isolates.

In this study, we present a near-chromosome level assembly of the *Bgh* reference isolate (DH14), which recovers approximately 30 Mb of previously unassembled sequence, supplemented with a new, manually curated annotation. Genome-wide comparative analysis of the European DH14 isolate with the Japanese isolate RACE1, which is the most divergent *Bgh* isolate sequenced so far (Lu et al. 2016), revealed clear evidence for large-scale conservation of gene order between isolates. Subsequent comparisons with genomes of closely related *B. graminis formae speciales* indicated extensive copy number variation (CNV) of genes encoding secretion signal-containing proteins (SPs), which mirrors the phylogenetic relationships of these host-specialized forms. Finally, we found evidence for recent proliferation of TEs in the *Bgh* genome and possibly other *formae speciales* of *B. graminis,* but not in powdery mildews colonizing dicotyledonous host plants. Collectively, these genomic features reveal unprecedented insights into *B. graminis* life history and co-evolutionary patterns of the fungal pathogen with grass hosts.

## RESULTS

### Large-scale *Bgh* genome organization

To facilitate a deep exploration of the *Bgh* genome, we applied third generation long-read DNA sequencing to generate high-quality genome assemblies of a European and a Japanese isolate, designated DH14 and RACE1. Whilst a short read-based genome is available for DH14 (Spanu et al. 2010), enabling direct comparison with the newly established long read-based assembly, isolate RACE1 was chosen because of its exceptionally high coding sequence divergence compared to a collection of 15 other *Bgh* isolates from different geographic origins, including DH14 (Lu et al. 2016). Although the PacBio platform-based sequence depth for isolate DH14 was relatively low (~25x; Table 1), the Canu (Koren et al. 2017) assembly resulted in 963 contigs (*i.e.* 14,093 fewer contigs than the published reference genome), a significant increase of the N50 statistic (now 4.6 Mb), and an almost complete recovery of previously unassembled genomic sequences (Table 1). Using existing data from sequenced plasmid and fosmid clones (Spanu et al. 2010), the assembly was further reduced to 318 scaffolds, comprising ~124.5 Mb in total. The final assembly was polished to remove erroneous base calls and insertions/deletions (indels) using short Illumina reads (~50x coverage). For isolate RACE1 the depth of the long read sequencing was higher (~50x) and thus these PacBio reads were used also for polishing. The resulting unscaffolded RACE1 assembly consists of 99 contigs (including the circular mitochondrial genome) and a total size of ~116.5 Mb (N50 3.9 Mb; Table 1). Overall, both assemblies show higher gene space coverage (BUSCO analysis) compared to the existing *Bgh* reference genome (Spanu et al. 2010) although the difference is comparatively small (Table S1). We did not find any evidence for the presence of previously reported *Bgh*-specific plasmid-like linear extrachromosomal DNA (Giese et al. 1990) in the two isolates.

**Table 1:**
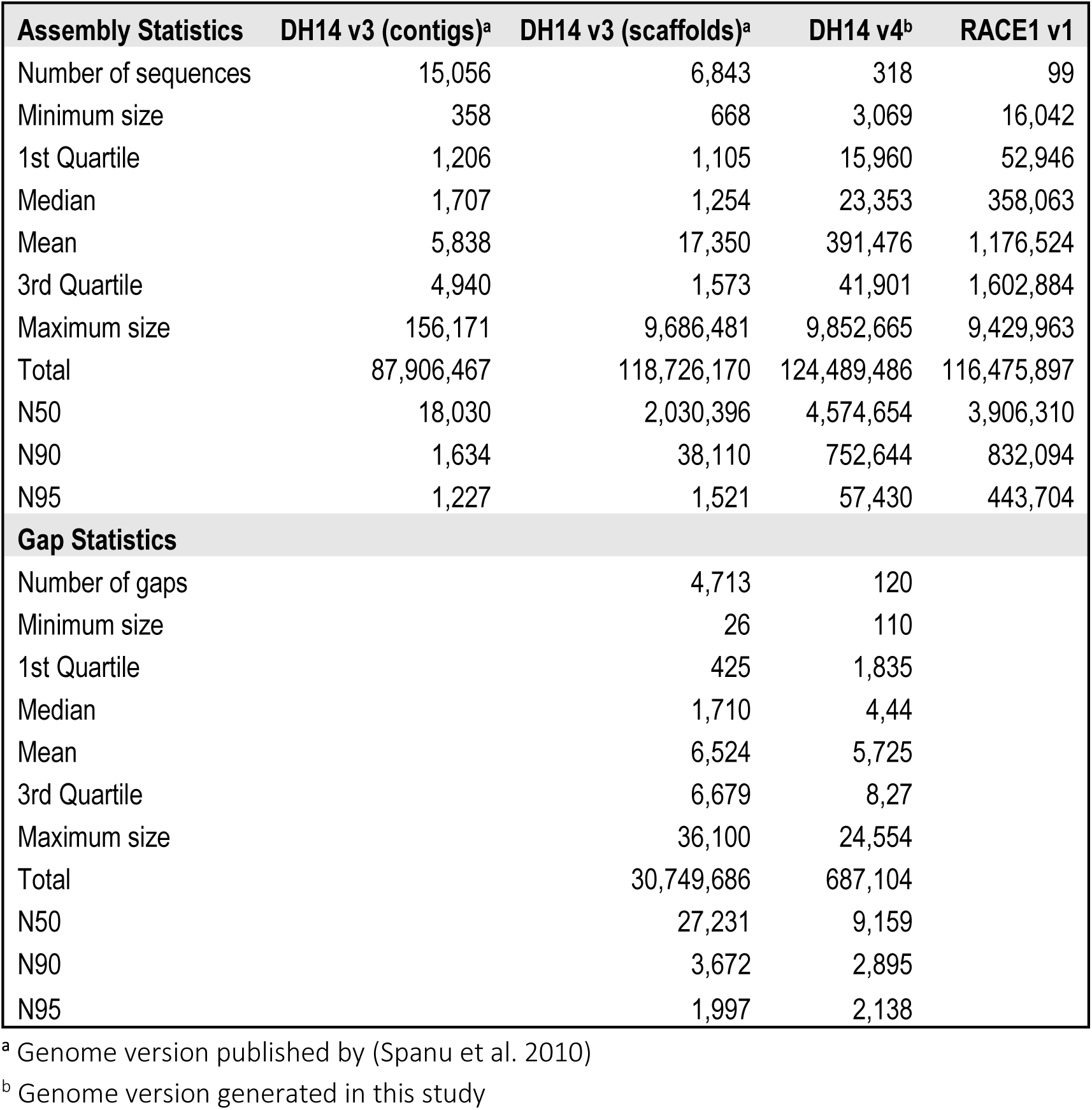
Assembly statistics for the genomes of the *Bgh* isolates DH14 and RACE1.

| Assembly Statistics | DH14 v3 (contigs)<sup>a</sup> | DH14 v3 (scaffolds)<sup>a</sup> | DH14 v4<sup>b</sup> | RACE1 v1 |
| --- | --- | --- | --- | --- |
| Number of sequences | 15,056 | 6,843 | 318 | 99 |
| Minimum size | 358 | 668 | 3,069 | 16,042 |
| 1st Quartile | 1,206 | 1,105 | 15,960 | 52,946 |
| Median | 1,707 | 1,254 | 23,353 | 358,063 |
| Mean | 5,838 | 17,350 | 391,476 | 1,176,524 |
| 3rd Quartile | 4,940 | 1,573 | 41,901 | 1,602,884 |
| Maximum size | 156,171 | 9,686,481 | 9,852,665 | 9,429,963 |
| Total | 87,906,467 | 118,726,170 | 124,489,486 | 116,475,897 |
| N50 | 18,030 | 2,030,396 | 4,574,654 | 3,906,310 |
| N90 | 1,634 | 38,110 | 752,644 | 832,094 |
| N95 | 1,227 | 1,521 | 57,430 | 443,704 |
| **Gap Statistics** |  |  |  |  |
| Number of gaps |  | 4,713 | 120 |  |
| Minimum size |  | 26 | 110 |  |
| 1st Quartile |  | 425 | 1,835 |  |
| Median |  | 1,710 | 4,44 |  |
| Mean |  | 6,524 | 5,725 |  |
| 3rd Quartile |  | 6,679 | 8,27 |  |
| Maximum size |  | 36,100 | 24,554 |  |
| Total |  | 30,749,686 | 687,104 |  |
| N50 |  | 27,231 | 9,159 |  |
| N90 |  | 3,672 | 2,895 |  |
| N95 |  | 1,997 | 2,138 |  |

<sup>a</sup> Genome version published by (Spanu et al. 2010)
<sup>b</sup> Genome version generated in this study

To further assess assembly quality and to facilitate future genome-anchored genetic studies, we compared the assemblies with a previously generated genetic map for *Bgh* (Pedersen et al. 2002). We located genomic positions for 80 mapped single-copy expressed sequence tag (EST) markers and compared their physical linkage with the corresponding genetic map. This revealed in most cases (67 out of 80 ESTs) a collinear marker order on physical and genetic maps (Suppl. Figure 1). In all but two cases in which discrepancies were found between physical and genetic maps, we observed collinearity of EST markers between the independently assembled genomes of DH14 and RACE1. Even large genetic linkage groups were mostly covered by only one or two genome contigs (e.g. linkage groups 2 to 7; Suppl. Figure 1), suggesting that our physical maps partly represent *Bgh* chromosomes or chromosome arms. In support of this, we identified 19 (DH14) and 20 (RACE1) cases of canonical telomeric repeat sequences (5’-TTAGGG-3’ hexamer; 34 to 61 tandem copies) at the beginning/end of contigs, in some instances together with distally-located gene-scarce regions likely resembling centromeres (Suppl. Figure 2, Table S2). The gene-scarce regions are in all cases associated with specific long interspersed nuclear elements (LINEs) of the *Tad1* family (Suppl. Figure 2).

The circular mitochondrial genomes of both isolates were closed, yielding total sizes of 104 kb (DH14) and 139 kb (RACE1) (Suppl. Figure 3A), which is in agreement with older experimental estimates (Borbye et al. 1992). Nucleotide sequence alignment indicated >96% identity of the mitochondrial DNA (mtDNA) of the two genomes. It further revealed that the RACE1 mitochondrial genome contains a ~32 kb duplication, while the DH14 mtDNA encompasses an ~1 kb isolate-specific sequence stretch that includes one predicted open reading frame (Suppl. Figure 3B). The structural and nucleotide differences might be linked to the isogamous and hermaphroditic manner of mitochondrial inheritance in *B. graminis* (Robinson et al. 2002), meaning that the mtDNA can originate from two parents derived from distant *Bgh* populations. Nonetheless, the *Bgh* mitochondrial genome does not present major differences in gene repertoires compared to known mitochondrial genomes of other Leotiomycetes (Mardanov et al. 2014), except for *Atp9,* which encodes the subunit 9/c of the mitochondrial ATP synthase complex and has been transferred to the nuclear *Bgh* genome. Consistent with this, *Bgh Atp9* carries an N-terminal mitochondrial transfer signal sequence. This gene translocation has been observed in other fungal species and might be related to a physiological adaptation, enabling transcriptional modulation of its expression in cell- and tissue-specific contexts (Bietenhader et al. 2012; Déquard-Chablat et al. 2011).

### Identification of isolate-specific genes, gene duplications and gene expression

Existing *Bgh* gene models for the isolate DH14 were transferred to the new assembly and were supplemented by new predictions generated by MAKER (Campbell et al. 2014), which were guided by protein and/or transcript evidence (whole-transcriptome shotgun sequencing; RNA-seq; see Materials and Methods). For RACE1, for which a prior genome annotation was unavailable, we generated *de novo* gene models using MAKER, guided by protein and transcript evidence from both *Bgh* isolates. We manually curated all gene models, removed poorly supported predictions, presumptive pseudogenes (mostly related to *Sgk2* kinase-like genes; [Kusch et al. 2014]) and annotations that overlapped with TEs. During the manual curation we noted several instances of (tandem) duplicated genes, which are highly sequence-related at the nucleotide level, and thus had been collapsed into single gene models in the previous DH14 genome assembly (Spanu et al. 2010). This complicated the annotation and therefore new gene identification numbers (IDs) were generated also for DH14 (Table S3).

The new annotation resulted in 7,118 gene models for DH14, of which 805 genes encode predicted SPs. A similar number of 7,239 gene models were found for RACE1 upon manual curation, including 770 that encode predicted SPs. A subgroup of SPs, called CSEPs, are secreted candidate virulence proteins defined by specific criteria (Pedersen et al. 2012). For a more comprehensive coverage of the deduced fungal secretome, we generally included all SPs in our analyses. This also allowed us to incorporate newly detected effector candidates resulting from the re-annotation of the *Bgh* genome. To compare the gene repertoires encoded by the DH14 and RACE1 isolates, we first used OrthoFinder to infer orthologous gene groups (orthogroups). This analysis identified 6,039 single-copy groups containing gene pairs with unambiguous one-to-one relationship between the isolates (Figure 1A). By manually incorporating additional position and synteny information from a whole-genome alignment (see below) for the inference of orthologous gene pairs, we could further resolve some ambiguities and identify additional relationships for unassigned genes with more dissimilar sequences, significantly increasing the number of one-to-one gene pairs to 6,844 (Figure 1B, Table S4). A comparison of DH14 and RACE1 orthogroups showed that most groups (6,200 out of 6,319) contain the same number of members in both isolates, but there are several groups with an isolate-specific expansion, indicating the presence of additional paralogs in one of the isolates (Figure 1C, Table S4). Such isolate-specific expansions occur almost 10-fold more frequently for SP-containing groups than for groups without SPs (9.4% and 1.2%, respectively; χ^2^-test, p<2e^-16^). An example for the occurrence of such an isolate-specific gene duplication is the *AVR_a1_* avirulence effector (Lu et al. 2016) for which two identical copies exist in DH14, while only one copy was found in RACE1 (Suppl. Figure 4A). By contrast, the *AVR_a13_* avirulence effector locus is highly similar in both isolates, with a single copy of *AVR_a13_* flanked by the other two members of the previously identified *AVR_a13_ CSEP* family (Lu et al. 2016; Pedersen et al. 2012) (Suppl. Figure 4B).

**Figure 1:**
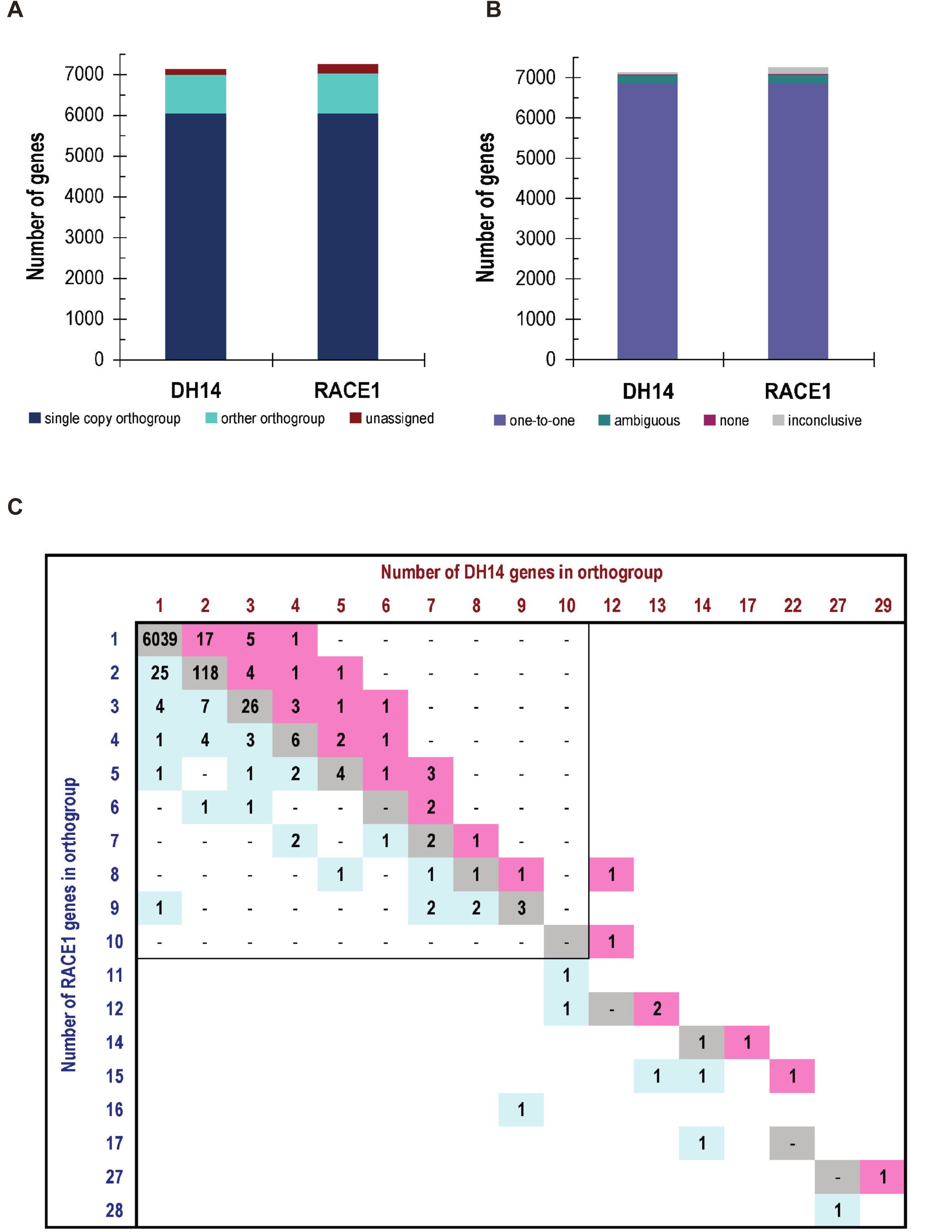
Identification of orthologous gene groups and gene pairs between the *Bgh* isolates DH14 and RACE1. A) Bar graph visualizing the number of gene models in DH14 and RACE1 that were assigned by OrthoFinder into single copy orthogroups (with one member per isolate), orthogroups with more than two members, or with no ortholog at all. (B) Bar graph summarizing the observed orthology relationships between DH14 and RACE1 gene models, as inferred from a combination of OrthoFinder results and additional manual inspection of gene positions and synteny. With this method, a one-to-one relationship between isolates could be established for most genes (6,844), while for ~200 genes in each isolate the relationship remained ambiguous (e.g. due to the existence of additional identical copies). Through further integration of RNA-seq data for both isolates, 31 and 27 genes were verified to be isolate-specific in DH14 and RACE1, respectively. For the remaining genes (47 in DH14 and 162 in RACE1) the relationship assignment was inconclusive due to still existing inaccuracies in the assemblies or annotations. (C) Graphical representation showing the composition of the identified orthologous gene groups from the OrthoFinder analysis. Most groups (6,200 out of 6,319) contain an equal number of members in DH14 and RACE1 (gray squares), while for some an isolate-specific enlargement can be observed with more members in one isolate than the other (light blue and pink squares).

We also searched for isolate-specific genes without any related sequence in the other respective isolate. For this purpose, we included the previously published RNA-seq data for RACE1 (Lu et al. 2016) and a corresponding newly generated dataset for DH14 as additional evidence to extract a high-confidence set of isolate-specific genes. This analysis identified in total 31 isolate-specific genes in DH14, for 13 of which we detected credible gene expression (FPKM (fragments per kilobase [sequence ength] and million [sequenced fragments])≥5) during pathogenesis (Table S5). A similar number of 27 isolate-specific genes was found in RACE1, of which 19 were also expressed (FPKM≥5) during pathogenesis (Table S5). Among these expressed isolate-specific genes, we found eight *SPs* in DH14, but only three in RACE1. As the two isolates are of opposite mating types, also the corresponding *MAT* isomorphs were among the isolate-specific genes, with DH14 carrying *MAT1-2-1* and RACE1 carrying both *MAT-1-1-1* and *MAT-1-1-3* (Table S5, Suppl. Figure 5).

Apart from a validation of the presence of isolate-specific genes, the RNA-seq data enabled us to examine also potential isolate-specific gene expression during infection. We searched for orthologous gene pairs for which we could detect robust transcript levels (FPKM≥10) in one of the two isolates while in the other the corresponding gene was not expressed (FPKM<1 or raw count≤2), and for which isolate-specific expression could be further validated by visual inspection in the Integrative Genomics Viewer (IGV; [Robinson et al. 2011]). A total of 15 genes showed differential expression based on these criteria (Table S6). Of these genes, 12 were specifically expressed in RACE1 (of which seven encode CSEPs), while three were expressed specifically in DH14 (Table S6), indicating substantial differences in expressed gene repertoires between *Bgh* strains.

### Genome synteny, structural and sequence variation between isolates

For a detailed genomic comparison, we conducted a whole-genome alignment of DH14 and RACE1 assemblies using MUMmer (Kurtz et al. 2004). Although RACE1 was chosen for genome sequencing based on its high sequence divergence to DH14 within coding regions (Lu et al. 2016), the genomes of the two isolates overall are still remarkably similar, with 92% and 98% of the assemblies of DH14 and RACE1 aligning to the corresponding other isolate at an average nucleotide sequence identity of ~99%. Moreover, the aligned sequence stretches form large syntenic blocks of up to 9 Mb, implying that gene order within the assembled contigs is also largely conserved between the isolates (Figure 2A). A closer inspection of the syntenic blocks revealed that the large-scale synteny between DH14 and RACE1 can be interrupted locally by intermittent stretches of non-syntenic alignments (e.g. to a different contig in the other isolate) or by sequence areas without a close match in the other genome (Figure 2B). These interspersed alignment gaps typically are rather small (<1 kb on average) and concern primarily regions of repetitive sequence, while only rarely affecting protein-coding genes (only 1% of alignment gaps affect genes). As both genomes are not resolved entirely to whole-chromosome level, we cannot estimate the full extent of large-scale chromosomal reshuffling. Nevertheless, the occurrence of within-contig alignment breaks provides evidence for at least two large-scale genomic rearrangements that involve genome stretches larger than 1 Mb (Figure 2A, Suppl. Figure 6). Additionally, we found 128 cases of genomic rearrangements within contigs, where sequence stretches of at least 10 kb are inverted relative to the other isolate. These inversions occur dispersed throughout the genome and the average size of inverted regions is around 20 kb with only seven regions larger than 50 kb. Roughly half of these local inversions (69 out of 128) affect gene-containing regions, but only for 22 of the corresponding regions we could verify by manual screening that they coincide with an inverted gene order relative to the flanking genes (Figure 2C, Table S4). In three of these cases, a further re-shuffling of genes was observed within the inverted region. Collectively, while large parts of the genome structure and gene order seem to be well conserved, we detect a number of mostly smaller synteny breaks that are dispersed throughout the genome and contribute to the structural variation between the two isolates.

**Figure 2:**
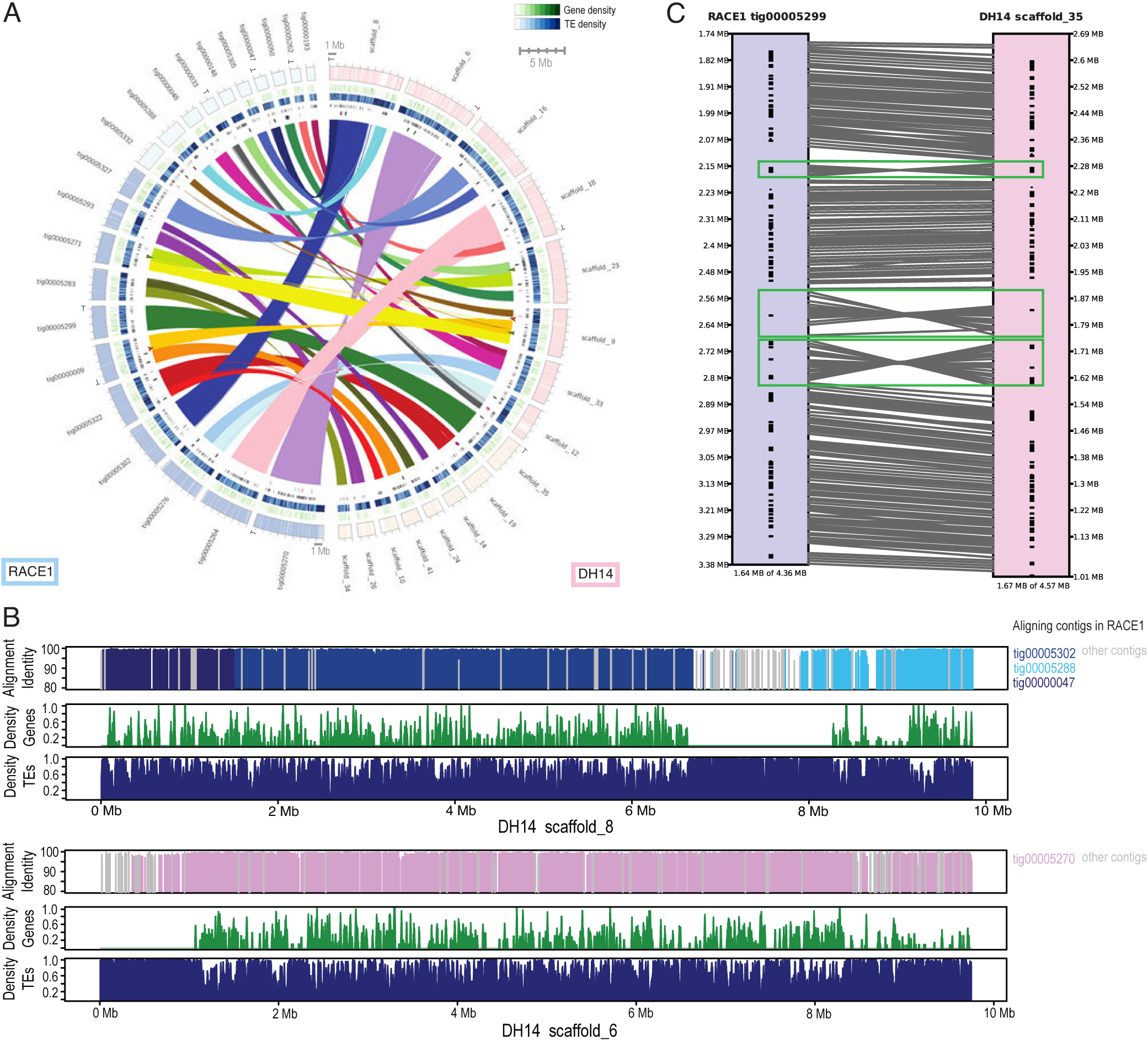
Syntenic relationships and genomic rearrangements between the genomes of *Bgh* isolates DH14 and RACE1. (A) Circos diagram illustrating the syntenic relationships between the genomes of DH14 and RACE1. Based on the corresponding N50 values, the 8 and 11 largest scaffolds/contigs from the DH14 and RACE1 genomes and the corresponding aligning scaffolds/contigs were chosen for visualization. Syntenic regions were identified based on whole-genome alignment, and aligning regions (>2 kb, similarity >75%) are connected with lines. The surrounding circles represent from the outside: on the right DH14 scaffolds (pink) and on the left RACE1 contigs (blue), with all unaligned regions (> 1 kb) indicated as white gaps; gene density (green) and TE density (blue) calculated in 10 kb sliding windows; locations of all genes predicted to code for SPs; locations of isolate-specific genes coding for SPs (dark red) or any other proteins (black) and isolate-specific additional gene copies/paralogs coding for SPs (pink) or any other proteins (grey). The arrowheads indicate observed alignment breaks of at least two large-scale genomic rearrangements (green and brown arrowheads, respectively). Scaffolds or contigs marked with "T" indicate presence of telomeric repeats. (B) Visualization of sequence identity to RACE1, gene and TE density along the two largest DH14 scaffolds. The top panel shows the sequence identity to the aligning RACE1 contigs obtained from the whole-genome alignment, with bars colored according to the involved RACE1 contig as indicated next to the graph. The two lower panels show the gene density (green) and TE density (blue) in DH14 calculated in 10 kb sliding windows along the DH14 scaffolds. (C) Detailed synteny between RACE1 tig00005299 and DH14 scaffold_35 visualized with SyMap (Soderlund et al. 2011). Local inversions within the otherwise syntenic region are highlighted by green boxes, while grey lines connecting the RACE1 and DH14 contigs represent the syntenic alignments and black marks on the contigs indicate the positions of annotated genes.

To examine the sequence variation between RACE1 and DH14, we used the single nucleotide polymorphisms (SNPs) identified in the MUMmer (Kurtz et al. 2004) dnadiff analysis to calculate SNP frequency in 10 kb sliding windows. In addition, we obtained SNPs for isolates A6 and K1 (Hacquard et al. 2013) from a short read alignment (see Materials and Methods) and calculated the corresponding SNP frequencies to re-examine the sequence variation of these isolates to the improved DH14 reference genome. On average, the overall SNP frequency is three times higher in RACE1 (4.7 SNPs/kb) than in A6 (1.4 SNPs/kb) and K1 (1.3 SNPs/kb). Moreover, a comparison of SNP frequency distributions between the three isolates shows that in RACE1 SNP frequencies below one SNP per kb are seen only rarely, whereas in A6 and K1 they are common (Suppl. Figure 7A). Accordingly, a two-component mixture model fitted to the observed SNP frequencies recovered the previously described (Hacquard et al. 2013) distinction between low and high SNP densities (mean ± standard deviation) for A6 (low: 0.1 ± 0.1; high: 1.9 ± 1.5) and K1 (low: 0.1 ± 0.1; high: 2.0 ± 1.7). By contrast, for RACE1 no such distinction could be observed and the SNP frequency was high for both model components (low: 3.8 ± 2.4; high: 10.6 ± 6.1; Suppl. Figure 7B).

### SP paralogs typically reside in close proximity

Although local clustering – in part even as tandem duplicates - of genes encoding effector candidates in the *Bgh* genome has been suggested and described (Pedersen et al. 2012), its scale at a genome-wide level remained unclear. This is mainly due to the severe fragmentation of the previously available genomic assemblies and the collapse of highly similar gene copies in the short read-based assemblies (Hacquard et al. 2013; Spanu et al. 2010). We therefore explored systematically whether *SPs* in general co-occur in close distance. Here we defined *SP* clusters based on two criteria: (1) each cluster contains at least three *SPs* and, and (2), two *SPs* are separated by a maximum of ten genes coding for non-secreted proteins. By these criteria, 72% of the SP-coding genes (583 out of 805) can be placed in three large clusters with more than 30 *SPs* and 74 smaller clusters with less than 20 *SPs* (Table S7). Consistent with an earlier study (Pedersen et al. 2012), many of these clusters harbor sequence-related genes which belong to the same orthogroup (Table S7), suggesting that they might originate from recent local duplications with subsequent sequence diversification, thus likely representing paralogs. Despite this occurrence of *SP* clusters, we did not observe local enrichment of *SPs* on particular genomic scaffolds (Suppl. Figure 8A). Rather we found that the *SP* count follows the scaffold size (Suppl. Figure 8B), which is in line with the results of a χ^2^-test that did not detect a significant deviation between the *SP* frequency per scaffold and the underlying genome fraction per scaffold (p=0.21).

### Copy number variation of *SPs* within and between *formae speciales* correlates with phylogeny and host specialization

To investigate the extent of within-genome gene duplications we used MCScanX (Wang et al. 2013) on the DH14 and RACE1 isolate datasets. As expected for a haploid genome, the majority of the genes exist in single copies, but ~10% have one or more paralogs (Table S8). Approximately one third of these duplications occur in tandem (36%), while the remaining ones are either proximal (in-between the next five genes) or dispersed throughout the genome (30% and 33%, respectively). When compared to the genomes of the phylogenetically closely related phytopathogenic fungi *Botrytis cinerea* and *Sclerotinia sclerotiorum,* the *Bgh* genome shows a higher percentage of duplications (11% *versus* 0.3% and 5.4%, respectively). A closer look at the *S. sclerotiorum* dataset revealed that the seemingly elevated number of dispersed and proximal duplications in this species is mainly comprised of retrotransposases that are retained in the corresponding annotation (Table S9). This finding indicates that the comparatively high number of paralogous gene pairs (812 out of 7118) in the *Bgh* genome is a unique characteristic among the so-far-sequenced Leotiomycetes.

We investigated whether these duplications can be associated with certain types of genes or functional domains and found that *SP* genes are significantly more subject to duplication than genes encoding non-SPs (χ^2^ test, p<0.001). Most duplications of *SPs* seem to occur in tandem (Table S10). Functional domain associations are poor for the group of *SP* genes because effector proteins often have few or no functional descriptions (applies to ~79% of the 805 predicted SPs in DH14 in PFAM-based searches; Table S11). However, there are two clusters with tandemly duplicated genes that are rich in genes encoding ribonuclease-like domains (SUPERFAMILY SSF53933, clusters 21 and 1), and two clusters with *Egh16* virulence factor homologs (PFAM PF11327, clusters 56 and 14). Among the genes coding for non-SPs, a portion of the duplications (181 out of 546) are related to genes with kinase-like domains (SSF56112, PS50011), which have been described previously as an over-proliferating family in the *Bgh* genome (Kusch et al. 2014).

Based on the above results we sought to determine whether gene copy numbers vary between strains belonging to different *formae speciales* of *B. graminis.* Using published datasets (Hacquard et al. 2013; Menardo et al. 2016; Menardo, Wicker et al. 2017; Wicker et al. 2013), we estimated the copy number of each *SP* based on the observed coverage of short read sequence alignments against the DH14 assembly (Figure 3A). To assess the accuracy of this analysis, a sample of genes with tubulin or actin functional domains and some additional non-SP-coding genes with conserved domains was used. As expected, this control dataset showed minimal variation and revealed conserved single-copy presence in all 52 genomes examined (representing 9 *formae speciales;* Suppl. Figure 9), indicating that the analysis based on coverage depth is robust. Nonetheless, false approximations cannot be fully excluded by this approach.

For the majority of *SPs* (458 of 805; 57%) we detected simple presence/absence variation between the different *formae speciales* and their respective isolates (Figure 3A). For a smaller fraction of *SPs* (201 of 805; 25%) the number of observed copies varies between 0 (absence) and more than 2 copies per genome. Interestingly, while variation in copy numbers between the examined genomes can be observed (Figure 3A), for many *SPs* (72%-87%, depending on the *forma specialis)* the number of gene copies is conserved among different isolates of the same *forma specialis* (e.g. BLGH_01048 has 2 copies in all f.sp. *secalis* and f.sp. *dicocci* genomes). In addition, high-copy *SPs* have low sequence similarity with each other (Figure 3B). To investigate whether CNV correlates with the phylogeny of the *formae speciales,* we generated a tanglegram using a dendrogram derived from the hierarchical clustering of the CNV data and a cladogram derived from a UPGMA tree based on ~1.07 million single nucleotide polymorphism (SNP) positions between the isolates (Figure 3C). The CNV-based dendrogram accurately groups the isolates based on their host specificity, indicating that isolates belonging to the same *formae speciales* have distinctive CNV profiles.

**Figure 3:**
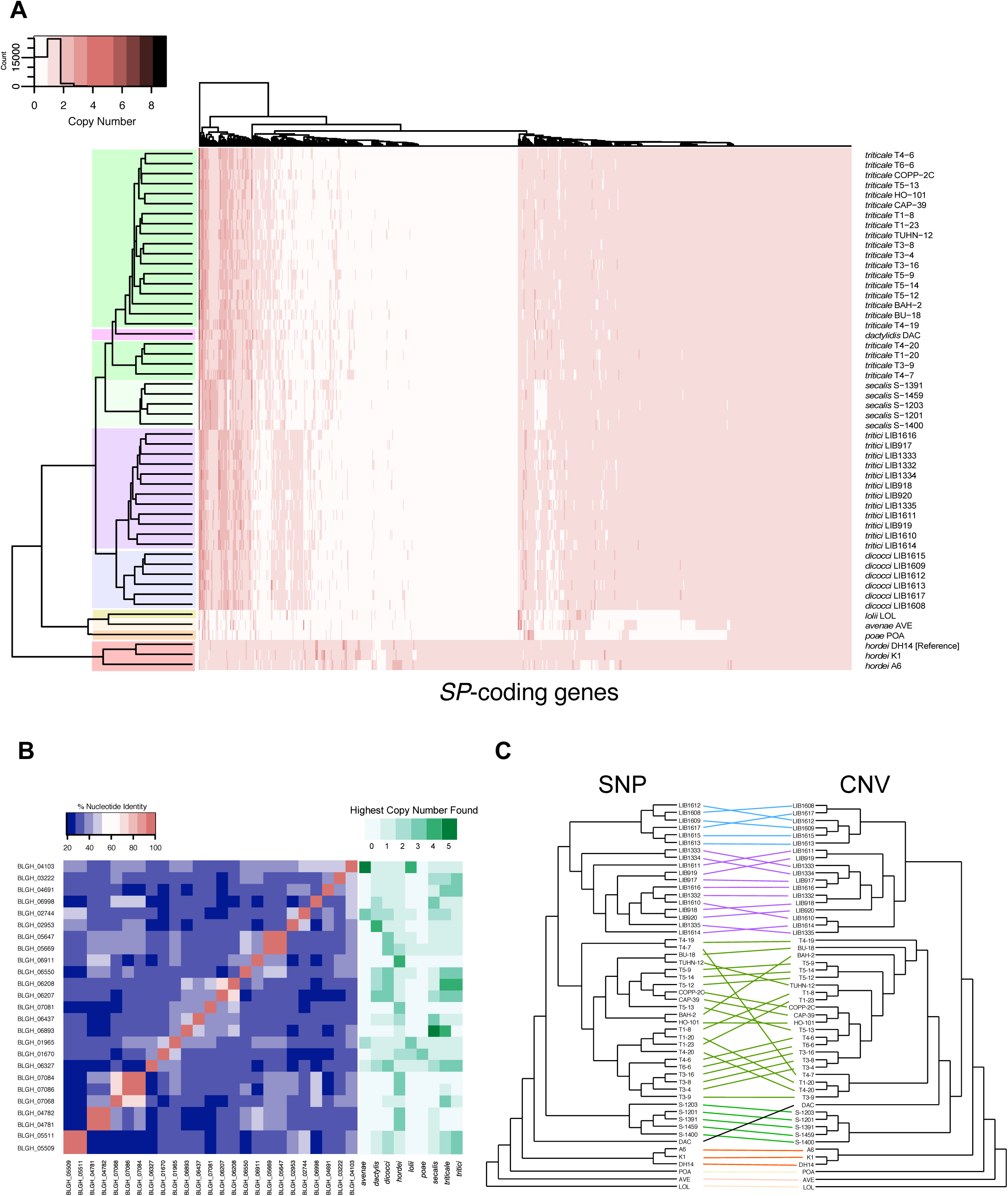
CNV of *SPs* reflects the phylogeny of various *formae speciales* and their isolates. (A) Heatmap illustrating the *SP* copy number per individual genome of various *B. graminis formae speciales (avenae, dactylis, dicocci, hordei, lolii, poae, secalis, triticale* and *tritici)* based on read mapping depth. Hierarchical clustering (Euclidean method) of the *SPs* and the isolates based on the CNV values is shown on the left and above the figure, respectively. Genomes belonging to the same *forma specialis* are color-highlighted on the left dendrogram. (B) Nucleotide identity matrix of high copy number *SP* subset (>3 in at least one isolate, excluding BLGH_02064, BLGH_02719, BLGH_07048 due to ambiguities in the gene model) resulting by alignments with EMBOSS-Needle (Rice et al. 2000). On the right, the highest copy number found in each *forma specialis* for these *SPs* is depicted. (C) Comparison between the hierarchical clustering dendrogram derived from the CNV analysis (Euclidean method) and the SNP-based UPGMA cladogram generated with SplitsTree (Huson and Bryant 2006) using a tanglegram generated with Dendroscope (Huson et al. 2007). Lines connect the same isolate, while colors correspond to different *formae speciales* (as in A).

### The *Blumeria* core effectorome

To define the core effectorome of the species *B. graminis,* we *de* novo-assembled and annotated the genomes of single isolates of the 9 *formae speciales* and inferred orthology relationships for the predicted proteomes. Our amino acid sequence-based orthology clustering of the predicted *SPs* (Suppl. Figure 10A) suggests that although part (see below) of the secretome is highly conserved in all *formae speciales,* another subgroup is divergent. Also, due to the divergence at the DNA sequence level the presence of certain *SPs* in the genomes of the more distantly related *formae speciales avenae, lolii* and *poae* was not detectable in the short read-based CNV analysis above (Figure 3A), while the orthology analysis identified related sequences at the amino acid level (Table S11). Yet, the *formae speciales avenae, lolii* and *poae* still share smaller intersections with the *Bgh* secretome compared to the rest (Suppl. Figure 10A).

Out of the 805 *Bgh SPs* present in isolate DH14, 442 have at least one ortholog in all genomes assayed. A considerable fraction of these widely conserved *SPs* (252 out of 442; 57%) has PFAM domains and/or homologs outside the *Blumeria* genus. As indicated by their functional annotation (e.g. peptidases/proteases, hydrolases), these SPs are rather part of a common SP repertoire of fungal plant pathogens and are not specific innovations of the grass powdery mildews. On the other hand, 190 SPs fulfil the typical CSEP criteria (no homology outside the Erysiphales, no PFAM domain; [Pedersen et al. 2012]) and can be considered as the core effectorome of the grass powdery mildews. These core CSEPs belong to different phylogenetic families (Suppl. Figure 10B), possibly targeting a core set of conserved host functions to maintain virulence on grasses.

### The *Bgh* genome exhibits no obvious compartmentalization

Various types of TEs that are dispersed more or less evenly throughout the genomes (Figure 4A, Suppl. Figure 2A) dominate the intergenic space of the DH14 and RACE1 genomes. Accordingly, many TEs can be found in close vicinity to genes, regardless if they are coding for SPs or not (Figure 4A). This pattern contrasts with other sequenced fungal and oomycete plant pathogens where transposon-rich areas are essentially limited to lineage-specific regions/chromosomes or are largely confined to isochores (Klosterman et al. 2011; Rouxel et al. 2011). Several copies of these elements seem to be expressed, in particular certain types of *Copia* elements (Table S12), and in many cases, overlap with the 5’ or 3’ UTRs of nearby genes (Figure 4A).

**Figure 4:**
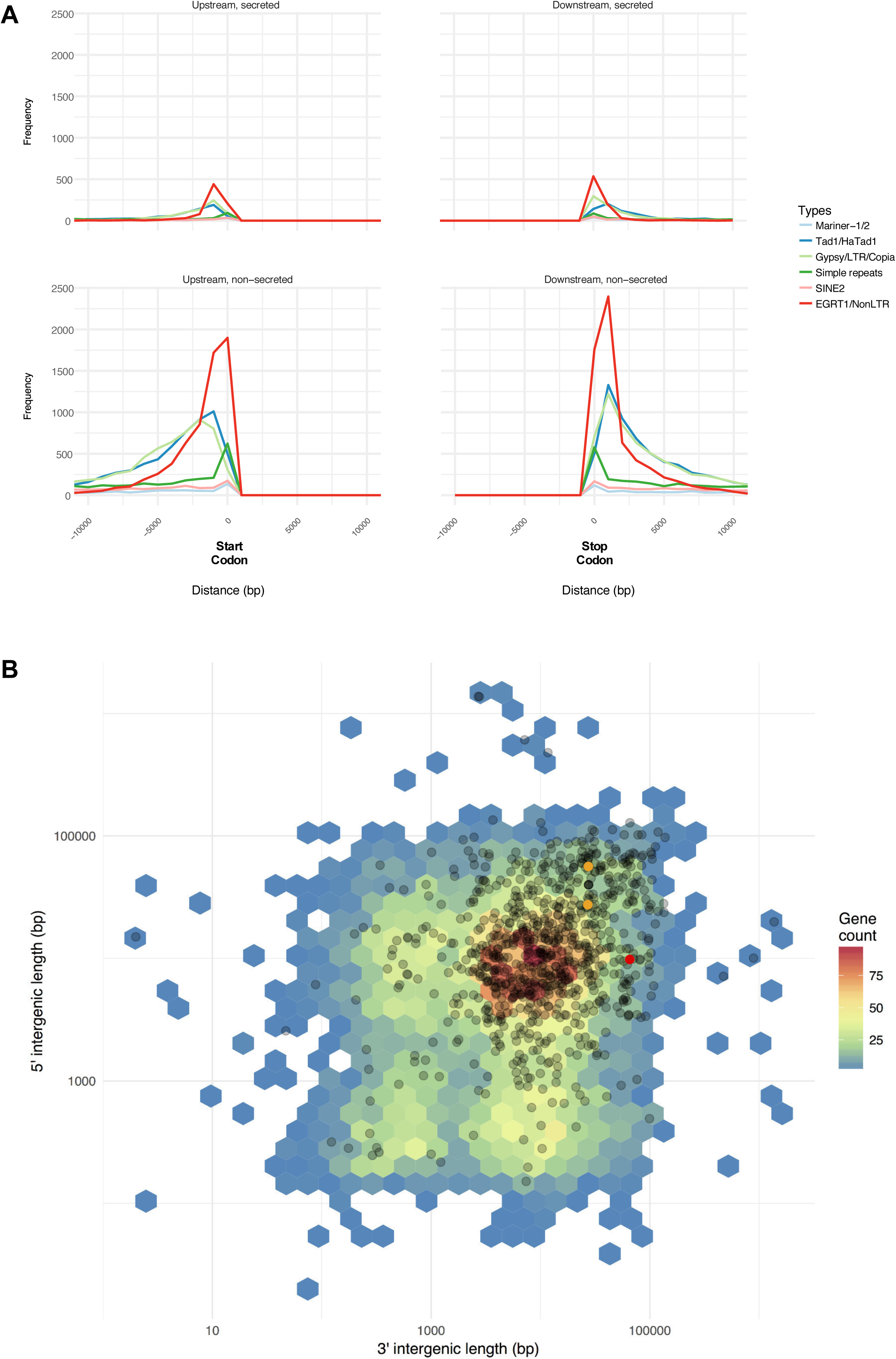
Intergenic space has similar size and is occupied by TEs in both *SPs* and non-SPs. (A) Frequency plot of the distance (−10,000 to +10,000 bp) of repetitive elements from the start codon (left panel) or the stop codon (right panel) of *SPs* (top panels) and non-SPs (bottom panels). The lines are color-coded and each represents a class of TEs according to the given legend. (B) 5’ and 3’ intergenic space size (y and x-axis) was calculated using BEDTools on the DH14 reference annotation. Black dots depict the intergenic length of all *SPs.* The orange dots mark the two *AVR_a1_* copies and the red dot marks *AVR_a13_*.

A complementary analysis of the local gene density, measured as flanking distances between neighboring genes, shows that in general the flanking distances in the *Bgh* genome are rather high, with an average distance of ~14 kb (Figure 4B). Accordingly, the surrounding genomic context of most genes in the *Bgh* genome is gene-sparse and repeat-rich and large flanking distances are not specific to *SP* genes (Figure 4B). In line with this pattern, also the flanking distances of the two known *Bgh AVR* effector genes, *AVR_a1_* and *AVR_a13_* (Lu et al. 2016), are not exceptionally large compared to the overall genome (Figure 4B). We further investigated whether genes coding for CSEPs or other SPs, which do not fulfil the typical effector criteria, present a difference in their 5’ or 3’ intergenic distances compared to ascomycete core ortholog genes *(COGs).* Regarding the 5’ intergenic distances, we detected no marked variation between the groups (ANOVA, p=0.382), while the 3’ intergenic distances on average were slightly larger for the *COGs* than for both the *CSEPs* and other *SPs* and (ANOVA, p=0.004; Tukey *post hoc* tests, p<0.05; Suppl. Figure 2B). The results of this analysis highlight that in *Bgh CSEPs* or other *SPs* are not located in peculiar gene-scarce regions compared to the conserved *COGs.* In addition, low gene density also could not be associated with high dN/dS rates (Suppl. Figure 2C), indicating that fast evolving genes in *Bgh* such as the *CSEPs* can occupy both gene-rich and gene-scarce areas. Thus, the *Bgh* genome does not appear to be split into distinct compartments, but is rather characterized by a low gene density and high TE density throughout the entire genome.

### A recent lineage-specific TE burst shaped the *Bgh* genome

Since TEs occupy the majority of the *Bgh* genome and are in many cases closely entangled with presumed virulence genes (SPs), we examined whether these repetitive sequences slowly accumulated over time or, alternatively, were subject to sudden expansions in the life history of powdery mildews, which might be linked to the observed proliferation and clustering of some highly sequence-related *SPs.* We used RepeatMasker (www.repeatmasker.org) to detect TEs with previously curated annotations found in Repbase (http://www.girinst.org/about/repbase.html), and subsequently generated repeat landscapes based on the divergence from the corresponding consensus TE sequences.

Surprisingly, most of the repetitive elements in *Bgh* show very low nucleotide sequence divergence (<10%) compared to the TEs in 13 closely related Leotiomycete genomes (typically 30%-40% nucleotide sequence divergence; Figure 5A, B), suggesting a recent lineage-specific expansion of TEs within *Bgh* (Figure 5B, C). In addition, there are 1,866 occurrences of long terminal repeats (LTRs) with less than 0.1% divergence associated with either *Gypsy* or *Copia* elements (~3% of the LTRs than can be identified), indicating that the *Bgh* genome carries very recent transposition events. Finally, the observed TE expansion in *Bgh* can be equally attributed to both LINE and LTR retrotransposons (Figure 5C). As outlined above, for part of these TEs, in particular *Copia* elements, evidence of expression can be found in the RNA-seq datasets (Table S12).

**Figure 5:**
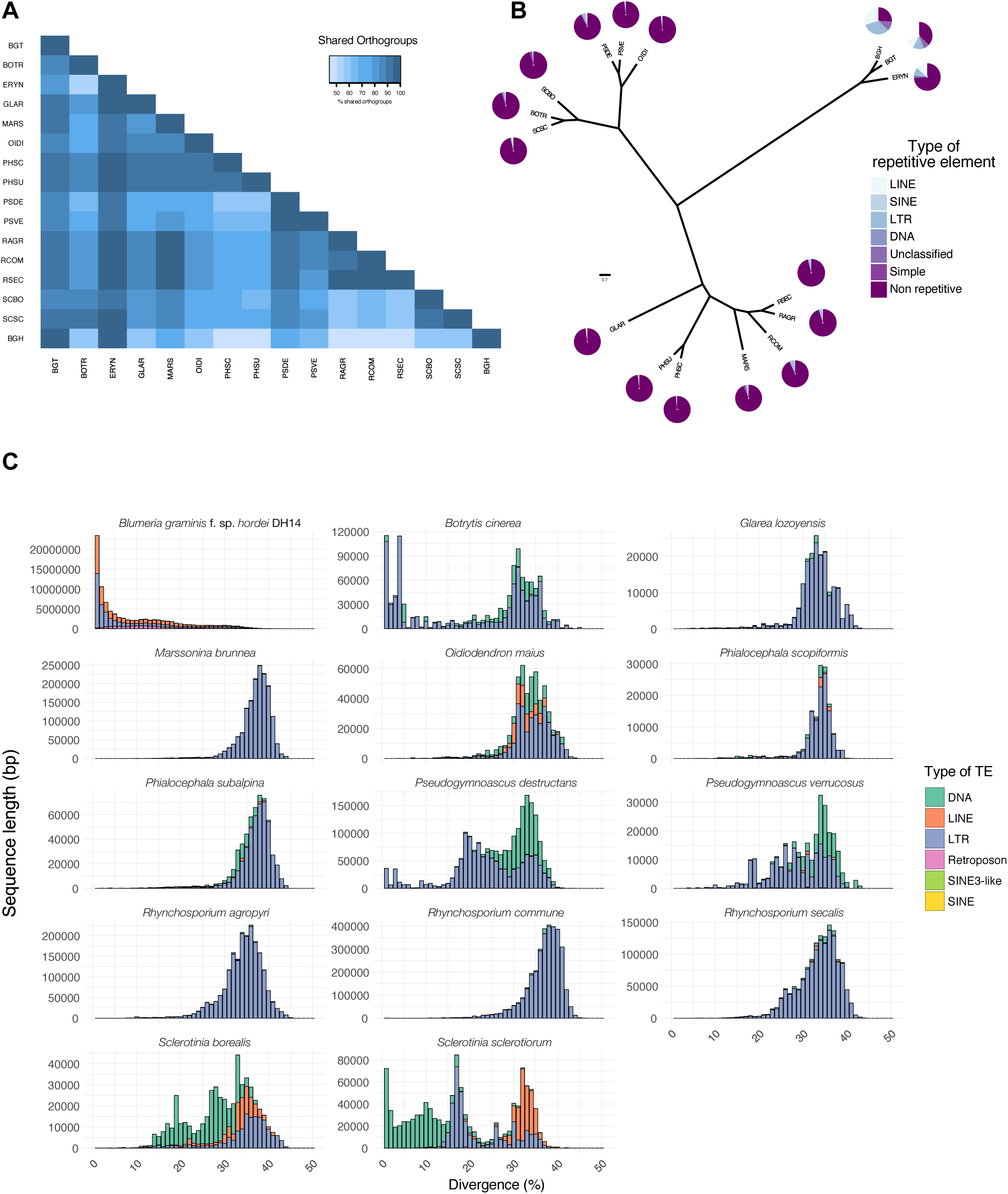
A recent TE burst in *Bgh.* (A) Orthogroup relationships of Leotimycete genomes. Leotiomycete genomes with available annotations in the NCBI database were assayed for their relatedness to the *Bgh* genome using OrthoFinder (Emms and Kelly 2015). Percentage of shared orthogroups is color-coded according to the key above (abbreviations of the species used can be found in Table S13). (B) Multi-locus phylogeny for 16 Leotiomycetes generated from 2,639 single copy orthologous genes identified with OrthoFinder (Emms and Kelly 2015) and drawn with FigTree (http://tree.bio.ed.ac.uk/software/figtree/). The repeat content, type and the portion it represents in each species’s genome is depicted in pie charts at the tip of each branch. The scale bar indicates the number of amino-acid substitutions per site. (C) Repeat landscapes for each genome of the tree shown in (B). Plots for *B. graminis* f.sp. *tritici* (BGT) and *E. necator* (ERYN) can be found in Suppl. Figure 9. The sequence length occupied in the genome is depicted on the y-axis (not normalized among the species examined), while the percent divergence from the corresponding consensus sequence is given on the x-axis.

Genome assemblies of *B. graminis* isolates belonging to other *formae speciales,* which are exclusively based on short reads, were found to underestimate both the magnitude of TE expansion and the presumed divergence time. This is due to the fact that the majority of the highly similar repetitive sequences collapse into few contigs, as revealed by the comparison of *Bgh* assemblies that are either based on long (PacBio) or short reads (Illumina; Suppl. Figure 11A). Therefore, for the other *formae speciales* of *B. graminis* it can only be assumed that they also experienced a recent TE expansion, while it remains unclear whether this event is older or more recent than the one in *Bgh.*

Remarkably, when applying the same pipeline on the sequenced genomes of the dicot-infecting powdery mildews *Erysiphe necator, E. pisi* and *Golovinomyces orontii,* the divergence from the consensus of the respective TE sequences is much higher (25%-35% compared to <10% in *Bgh),* suggesting that the expansion of the repetitive elements in these species is more ancient than in *Bgh* (Suppl. Figure 11B). This calculation is unlikely to be an underestimation due to the short read-based genome assemblies of these species as long (PacBio) or short read-based assemblies revealed similar divergence rates for *G. orontii.* Because evolutionary rates within the *Erysiphaceae* family appear to be comparable (Mori et al. 2000; Takamatsu and Matsuda 2004) and essentially all TEs in dicot-infecting powdery mildews are sequence-diverged (>10%; Suppl. Figure 11B), this observation suggests independent “transposon bursts” for each powdery mildew lineage that occurred at different times.

## DISCUSSION

### An improved assembly provides insights into large-scale organization of the *Bgh* genome

The genome of the obligate biotrophic pathogen *Bgh* is characterized by a loss of genes encoding enzymes of primary and secondary metabolism as well as an expansion of overall genome size due to a massive proliferation of TEs (Spanu et al. 2010). This high repeat content, with TEs representing more than two thirds of the genome, makes it essentially impossible to generate chromosome-level DNA assemblies from short sequencing reads that do not allow to resolve these highly similar sequences. Accordingly, the first short read-based assembly for *Bgh* isolate DH14 was highly fragmented, with more than 15,000 contigs (and close to 7,000 scaffolds), and about one third of the estimated genome size was not covered (Spanu et al. 2010), possibly due to collapsed sequences.

We here used a single-molecule sequencing technique to generate long DNA sequence reads, which enabled us to establish high quality genome assemblies for the two *Bgh* isolates, DH14 and RACE1. For DH14, this long-read-based assembly showed a >10-fold improved contiguity and recovered a substantial amount of previously unassembled genomic sequence (50% increase in genome size) compared to the first genome draft. In both DH14 and RACE1 assemblies, the largest contigs are more than 9 Mb in size, likely to represent complete chromosome arms, and a similar number of observed telomeric repeat regions in both assemblies (19 and 20) suggests the *Bgh* genome is likely partitioned into 10 chromosomes.

The high assembly quality is also supported by both an improved gene space coverage (now >98% BUSCO coverage for the newly annotated DH14 reference genome; Table S2) and good agreement with a previously published genetic map of *Bgh* (Pedersen et al. 2002). The yet missing BUSCOs could be due to either real gene loss events or failure to detect the corresponding conserved ortholog by the software, suggesting that the core gene space is now essentially completely covered in the *Bgh* reference genome. The few observed discrepancies between physical contigs and the genetic map might be attributed to the fact that the linkage map was constructed from a cross between two isolates (C15 and JEH31) that are different from the ones used in this work. Therefore, while we cannot exclude that the few discrepancies are at least partly due to remaining inaccuracies in either our assemblies or the genetic map, these also could be evidence for additional isolate-specific genomic rearrangements.

### Effector repertoires differ slightly between the two *Bgh* isolates

Approximately 74% of both assemblies are made up of repetitive elements that are uniformly dispersed across the genome, which is an even higher repeat fraction than predicted before for *Bgh* (64%; Spanu et al. 2010). The previous underestimation of the TE content could be due to the collapse of highly repetitive short read-based sequences during genome assembly. While a comparable fraction of TEs was described for the oomycete pathogen *P. infestans* (Haas et al. 2009), other sequenced fungal genomes contain markedly lower fractions (Figure 5B; [Castanera et al. 2016]).

In addition to a drastically improved assembly of repetitive sequences, we also noticed the existence of loci with similar or identical copies of a number of genes, which had previously been collapsed into single gene models (Table S3). Thus, genome re-annotation based on the new assemblies also provided an improved representation of *Bgh* gene repertoires, increasing the number of gene models from 6,470 (Spanu et al. 2010) to 7,118 in the annotation of the reference isolate DH14. A subsequent comparison of protein-coding genes between DH14 and RACE1 revealed largely conserved gene numbers between the two isolates, which is in agreement with previous observations based on short read-based assemblies of three *Bgh* isolates (Hacquard et al. 2013). However, due to the improved resolution, here we were able to identify several cases of isolate-specific gene family expansions and gene duplications, especially affecting *SPs.* Moreover, we identified several *SP* genes that were present exclusively in one of the genomes and lacked any similar sequence in the other isolate. The observed differences suggest that diversity of *SP* repertoires in *Bgh* is maintained mostly through gene duplications with subsequent sequence diversification and gene deletions. Thus, our observations for *Bgh* reflect the general evolutionary pressure on pathogen populations to diversify effector repertoires, which could then serve as reservoirs for rapid adaptation in response to population-level alterations in host *R* genes. Accordingly, the diversity of *Bgh* effectors is important in balancing the trade-off between ensuring virulence function and, at the same time, trying to escape detection by the host plant (Lo Presti et al. 2015).

Interestingly, the gene with the strongest isolate-specific expression in RACE1 encodes thioredoxin A, which is important for protection from oxidative stress and contributes to virulence of human pathogenic bacteria and fungi (Cheng et al. 2017; Cintra et al. 2017). However, as the RNA-seq samples for RACE1 and DH14 were generated in different experimental batches, we cannot fully rule out the possibility that the expression differences we observed could be partially influenced by batch effects.

### TEs and *SPs* are evenly dispersed throughout the genome

Many filamentous pathogens exhibit a distinct genome architecture, denoted as “two-speed genome”, with well-defined blocks of low gene and high TE density, interspersed between the generally more prevalent genomic areas of high gene and low repeat content (Dong et al. 2015; Faino et al. 2016; Jonge et al. 2013). These TE-rich blocks, which often harbor genes encoding secreted effector proteins, typically exhibit high lineage-specific diversity and are prone to be involved in genomic rearrangements (Faino et al. 2016; Jonge et al. 2013; Yoshida et al. 2016). In this way, these regions are thought to provide a pool of genetic variation that is needed by phytopathogens to quickly adapt to changing requirements in the evolutionary arms race with their hosts (Dong et al. 2015).

In *Bgh,* however, the situation is clearly different, as the numerous TEs are not restricted to specific areas, but rather evenly dispersed throughout the genome (Figure 2A, Suppl. Figure 8A). In addition, neither are the flanking regions of *SPs* particularly enriched in TEs, nor are they markedly larger compared to non-SPs (Figure 4A, B). Moreover, *SPs,* whether they are categorized as putative effectors (CSEPs) or not, are not associated with gene-sparse (Suppl. Figure 2B) or peculiar genomic regions (Suppl. Figure 8A), but their number is positively correlated with scaffold size (Suppl. Figure 8B). Additionally, the dN/dS ratio of the *CSEPs* is not associated to local gene density (Suppl. Figure 2C).

We also did not detect any large lineage-specific regions as reported for *Verticillium dahliae* (Jonge et al. 2013). Instead, smaller lineage-specific (<1 kb on average; up to 51 kb) or locally inverted (<20 kb on average; up to 90 kb) sequence stretches can be found dispersed rather evenly distributed throughout the genomes of the two *Bgh* isolates. Thus, the organization of the *Bgh* genome does not match the “two-speed genome” model (Dong et al. 2015), in which genetic variation is concentrated in specific genomic areas. Instead, *Bgh* appears to have a “one-speed/high-speed genome” where genetic and structural variation is not concentrated in specific compartments but rather sustained throughout the whole genome. Such a genome architecture might contribute to maintaining genetic diversity of mainly asexually reproducing *Bgh.* However, in this scenario genomes would be expected to rapidly lose the overall synteny due to TE activity and cumulative effects of local genome rearrangements. Thus, it is conceivable that occasional sexual reproduction ensures the maintenance of overall synteny of *Bgh* genomes.

Our present work supports the assumption of a predominantly asexual reproduction mode in *Bgh,* as we were able to recover the previously described mosaic genome structure in the European *Bgh* isolates (A6, K1 and DH14), with isolate-specific alternating regions of low and high sequence diversity (Hacquard et al. 2013) (Suppl. Figure 7). For the highly divergent Japanese isolate RACE1, on the other hand, no monomorphic regions were detectable relative to DH14 (Suppl. Figure 7), which is most likely due to the prolonged geographic separation of the two isolates during which sequence variation could accumulate at a whole-genome scale.

### Grass powdery mildews have a fast-paced secretome adapted to their respective hosts

Effector proteins play a crucial role in interactions between plant pathogens and their respective hosts (Wit et al. 2016), and consequently both small (sequence divergence) and big (loss of effector clusters) changes can drive the preference of the pathogen to a new host (Inoue et al. 2017). To date, several genome reports have established that phylogenetically related pathogens share a core effectorome, whereas each member of a taxonomic lineage contributes its own unique effectors to the pan-effectorome (Chiapello et al. 2015; Hartmann et al. 2017; Ma et al. 2010). The large number of candidate effectors in the core effectorome of the *Blumeria* genus identified here, including at least 190 CSEPs belonging to 74 gene families, suggests these are indispensable for the maintenance of fungal virulence on different monocotyledonous hosts in each *forma specialis* of the species *B. graminis*. Whether the corresponding effector families mainly target different host components belonging to few or a large number of cellular pathways for the establishment of a biotrophic relationship with their grass hosts remains to be tested.

One interesting aspect of the grass powdery mildew effectorome is the CNV that some of its members experience (Figure 3A, B). As in other plant pathogens, this variation is dominated by presence/absence polymorphisms (Hartmann et al. 2017), indicating strong selection for some *SPs* by certain host genotypes. In addition, we noted increased numbers of *SP* copies in particular isolates and *formae speciales,* suggesting that transcript dosage might also play a role in host adaptation of powdery mildews. In other plant pathogens, increased copy number of virulence genes can alter the infection phenotype, as for example reported in the case of *ToxB* in *Pyrenophora tritici-repentis* (Manning et al. 2013; Martinez et al. 2004).

For powdery mildews this evolutionary pattern might be particularly advantageous because the loss of the repeat-induced point mutation mechanism (RIP; [Spanu et al. 2010]) allows additional gene copies to remain intact and functional (Freitag et al. 2002), providing a presumed fitness advantage. This can be for example observed in *Eryshipe necator,* where an increased copy number of *EnCYP51* enhances fungicide resistance (Jones, L. et al. 2014). Similarly, careful re-examination of existing data using the information from the new *Bgh* reference assembly indicates that for some isolates duplications of *CSEPs* could offer the means to escape detection by their respective host *via* naturally accumulating mutations in one of the copies. An example of this might be *AVR_a1_*, where one of the two copies present in isolate CC107 has accumulated mutations allowing evasion of detection in barley cultivars carrying the matching *Mla1 R* gene (Lu et al. 2016). A recent report (Menardo, Praz et al. 2017) suggests extensive *forma specialis-specific* expansions of certain *CSEP* families, supporting the conclusions of our CNV and *SP* orthology analysis (Figure 3A, Suppl. Figure 10A).

### TEs expanded suddenly and massively in the *Erysiphaceae*

The *Bgh* genome is frequently referred to as a typical example of repeat-based expansion of an eukaryotic genome (Raffaele and Kamoun 2012). Even early studies predating high-throughput genome sequencing revealed that the effect of TEs in this pathogen’s genome is significant. This conclusion was based on the frequency these sequences are associated with coding regions (Oberhaensli et al. 2011; Rasmussen et al. 1993; Wei et al. 1996). Nevertheless, the question of whether the activity of TEs and their dominance in the genome has been a beneficial or a neutral feature is still open.

TEs in the genome of *Bgh* are evenly distributed, in part transcriptionally active and flank virulence genes as much as genes involved in all types of basic biological processes (Figure 2A, Suppl. Figure 2, Suppl. Figure 8A). As in many other cases, it can be hypothesized that TEs can act as templates for rearrangements, deletions and duplications of genomic sequences (Faino et al. 2016; Hartmann et al. 2017). Furthermore, TE insertions next to or within virulence genes can change the pathogen’s host range (Ali et al. 2014; Kang et al. 2001).

We show for the first time that TEs in the grass-infecting (*Blumeria*) and dicot-infecting (*Erysiphe*) powdery mildews experienced sudden and, in evolutionary terms, synchronous expansions. Taking into consideration molecular clock studies (Menardo, Wicker et al. 2017; Mori et al. 2000), it is tempting to hypothesize that TE bursts in the genomes of Erysiphaceae occurred independently of each other and might have preceded or followed adaptation to new hosts. Similar observations placing TE bursts around speciation times have been reported in the plant pathogen *Leptosphaeria maculans* (Grandaubert et al. 2014; Rouxel et al. 2011) and other eukaryotic organisms (Rebollo et al. 2010). Theoretical models suggest that sudden TE expansions, when seen as a source of mutations, can push asexual organisms to a fitness optimum in adverse conditions (McFadden and Knowles 1997; Startek et al. 2013). Given that at least powdery mildews of the genus *Blumeria* reproduce mainly clonally (asexual) as haploid organisms (Hacquard et al. 2013; Wicker et al. 2013) and their *formae speciales* exhibit narrow host specificity, our findings call for future studies to clarify the relationship between TE expansion and changes in the pathogen’s host range.

## CONCLUSIONS

We provide a greatly improved reference (isolate DH14) resource for the barley powdery mildew pathogen, and a near-continuous assembly of the highly divergent isolate RACE1. Gene order between these two isolates is retained at large scale, but locally disrupted. Using the new reference and supplementary transcriptomic and genomic data, we reassessed the secretome of grass powdery mildews and defined a core group of 190 SPs, which are likely to be indispensable for virulence. Inter-*formae speciales* comparisons further revealed that these virulence-related genes exhibit extensive CNV and sequence divergence, which reflects the phylogeny of these powdery mildews. *SP* genes are often locally clustered, but these clusters are evenly dispersed throughout the genome. TEs, which like the *SP* clusters are uniformly distributed in the *Bgh* genome and in part actively transcribed, experienced a recent lineage-specific expansion.

Taken together the results presented here indicate that *Bgh,* and more broadly the species *Blumeria graminis,* has a highly dynamic genome. While for other filamentous pathogens the existence of a “two-speed” genome has been suggested, the characteristics of the *Bgh* genome (even genome-wide distribution of TEs and *SPs*) indicate a “one/high-speed” genome for this pathogen and possibly its close relatives. It remains to be shown whether and how these features were enabled by the loss of genome defense modules (e.g. RIP), and if they contributed as springboard for the conquest of new host species (host jumps and host range expansions).

## MATERIALS AND METHODS

### Genome sequencing

For DH14, genomic DNA was extracted as described in (Spanu et al. 2010), while for RACE1 the protocol described in (Feehan et al. 2017) was used. Subsequently, SMRTbell™ genomic libraries were generated and sequenced at the Earlham Institute (formally known as The Genome Analysis Centre, Norwich, United Kingdom) and at the Max Planck Genome Centre in Cologne (Germany) for DH14 and RACE1, respectively. The Pacific Biosciences (PacBio) RSII sequencing platform with either P5C3 (DH14) or P6C4 (RACE1) chemistry was deployed (Pacific Biosciences, Menlo Park, CA; (Eid et al. 2009)). A total of 21 SMRT cells achieved ~50x coverage for RACE1, while for DH14 6 SMRT cells resulted in ~25x coverage. In addition, DH14 genomic DNA was sequenced at ~50x coverage with the Illumina MiSeq platform, providing 2x300 bp paired-end reads.

### Genome assembly

For both isolates the obtained PacBio reads were trimmed, corrected, and assembled using the Canu assembler (version 1.4; [Koren et al. 2017]) with default settings. The RACE1 assembly was further polished using Quiver (version 0.9.0; [Chin et al. 2013]) with default parameter settings. In the case of DH14 the resulting contigs were scaffolded with BESST (version 2.2.5; [Sahlin et al. 2014]) using previously published plasmid and fosmid libraries (Spanu et al. 2010) and then polished using Illumina short reads and Pilon (version 1.18; [Walker et al. 2014]). To assess completeness of both assemblies we applied BUSCO (version 2.0.1; [Simão et al. 2015]) with default parameters searching against the Ascomycota database (ascomycota_odb9). To compare the assemblies with a previously published genetic map for *Bgh* (Pedersen et al. 2002), we obtained the nucleotide sequences of 80 single copy EST markers from this study and used BLASTN (BLAST+ version 2.3.0; [Camacho et al. 2009]) to map these sequences against our genome assemblies (with –e-value 1e^-6^), thereby revealing their genomic location.

### RNA sequencing and alignment

For RACE1, we used RNA-seq data generated in the context of a previous study (Lu et al. 2016), and for DH14 we generated corresponding samples from barley leaf epidermal peels at 16 and 48 h after *Bgh* conidiospore inoculation for RNA-seq as described before (Lu et al. 2016). The RNA-seq libraries were prepared by the Max Planck Genome Centre in Cologne (Germany) using the Illumina TruSeq stranded RNA sample preparation kit. The resulting libraries were subjected to paired-end sequencing (150 bp reads) using the Illumina HiSeq2500 Sequencing System.

To assess gene expression in DH14 and RACE1, the RNA-seq reads from both isolates were mapped to both genome assemblies under consideration of exon-intron structures using the splice aware aligner TopHat2 (Kim et al. 2013) with adjusted settings (‐‐read-mismatches 10 ‐‐read-gap-length 10 ‐‐read-edit-dist 20 ‐‐read-realign-edit-dist 0 ‐‐mate-inner-dist 260 ‐‐mate-std-dev 260 ‐‐min-anchor 5 ‐‐splice-mismatches 2 ‐‐min-intron-length 30 ‐‐max-intron-length 10000 ‐‐max-insertion-length 20 ‐‐max-deletion-length 20 ‐‐num-threads 10 ‐‐max-multihits 10 ‐‐coverage-search ‐‐library-type fr-firststrand ‐‐segment-mismatches 3 ‐‐min-segment-intron 30 ‐‐max-segment-intron 10000 ‐‐min-coverage-intron 30 ‐‐max-coverage-intron 10000 ‐‐b2-very-sensitive) to account for sequence variability between isolates. To assess the expression of individual genes, we obtained raw fragment counts per gene from the mapped RNA-seq reads for both isolates (summarizing both time-points) using the featureCounts function (-t CDS -s 2 -M -p) of the Subread package (version 1.5.0-p1;(Liao et al. 2014)) and subsequently normalized these raw counts to fragment counts per kilobase CDS per million mapped reads (FPKM) for better comparability.

Expression of TEs in the isolate DH14 was assessed by mapping pooled RNA-seq reads coming from the 16 and 48 hpi DH14 samples with STAR (Dobin et al. 2013), using the RepeatMasker-derived gff file as annotation. Raw counts per TE annotation were obtained using the ‐‐quantMode GeneCounts option.

### Gene annotation

The prediction of DH14 and RACE1 gene models was performed using the MAKER pipeline (version 2.28; [Holt and Yandell 2011]), which integrates different *ab initio* gene prediction tools together with evidence from EST and protein alignments.

For DH14, initially the previous gene models (v3, www.blugen.org) were transferred to the new assembly as described (Campbell et al. 2014). Then an additional round of annotation followed, incorporating ESTs assembled from public *Bgh* datasets (Table S13) using Trinity (Haas et al. 2013), protein datasets (Table S13), as well as trained prediction models for AUGUSTUS (Stanke et al. 2006), SNAP (Korf 2004) and GeneMark-ES (Ter-Hovhannisyan et al. 2008) as supporting evidence. All the annotations were subsequently manually curated using Web Apollo (Lee et al. 2013), removing unsupported gene models.

For RACE1, we performed a complete *de novo* annotation, as there were no previous gene models available. For this purpose, the MAKER pipeline was first run using AUGUSTUS (Stanke et al. 2006) with species model *Botrytis cinerea* and GeneMark-ES (Ter-Hovhannisyan et al. 2008) for *ab initio* gene prediction together with transcript and protein alignment evidence. The corresponding alignment evidence was created from BLAST and Exonerate (Slater and Birney 2005) alignments of the DH14 protein sequences as well as RACE1 protein and transcript sequences. The RACE1 transcript and protein sequences for these alignments were obtained from the corresponding RNA-seq data *via* a transcriptome *de novo* assembly using Trinity (Haas et al. 2013); with default parameter settings for paired-end reads) and subsequent open reading frame/peptide prediction using TransDecoder (Haas et al. 2013); with default settings). The resulting gene models from the first MAKER run were used as initial training set for another *ab initio* prediction tool, SNAP (Korf 2004). Next, the annotation pipeline was re-run including all three *ab initio* prediction tools together with the transcript and protein alignment evidence, thus generating a second, improved training set for SNAP. After re-training SNAP on this set, the complete annotation pipeline was run a third time to yield the final RACE1 gene models. For both isolates, the obtained gene models were manually curated using Web Apollo (Lee et al. 2013), to correct for errors and remove poorly supported gene models. The mitochondrial genome of DH14 was annotated using RNAweasel and MFannot (http://megasun.bch.umontreal.ca/RNAweasel/).

### Identification of orthologous genes and gene groups

Groups of orthologous genes (orthogroups) were inferred from DH14 and RACE1 using OrthoFinder (version 1.1.8; [Emms and Kelly 2015]) with the inflation value I set to 1.2. To further resolve ambiguities in the orthogroups and detect additional relationships between more dissimilar sequences, subsequently, a manual screening of gene positions and co-linearity in the two genomes was performed and the ortholog assignment was refined accordingly.

Isolate-specific genes were identified by combining the results of the OrthoFinder analysis with the alignment results for the RNA-seq data from both isolates. Explicitly, a gene was only considered to be specific for one isolate if, after OrthoFinder analysis and manual refinement, there was no orthologous gene detectable in the genome of the other isolate and additionally also no RNA-seq fragment (read pair) from the other isolate were detected to map against this gene (raw count ≤1). The fragment count per gene was calculated from the mapped RNA-seq read pairs (with mapping quality >0) using featureCounts (version 1.5.0; [Liao et al. 2014]) with adjusted settings (-s 2 -p -M), based on the curated gene models. For identification of isolate-specific gene expression the raw fragment counts were further normalized to FPKM values, to adjust for potential differences in coding sequence length and RNA-seq read depth between isolates.

To calculate non-synonymous (dN) and synonymous (dS) substitution rates between DH14 and RACE1, we first aligned the protein sequences for each manually of the curated orthologous gene pairs with ClustalW (version 2.1; [Larkin et al. 2007]). Subsequently, the protein alignments were converted to codon alignments using PAL2NAL (version 14; [Suyama et al. 2006]) and dN and dS rates were estimated from these codon alignments using the yn00 function of the PAML package (version 4.4; [Yang 2007]).

### Whole-genome comparison

A whole-genome alignment between DH14 and RACE1 was generated using the nucmer and dnadiff functions of the MUMmer software (version 3.9.4; [Kurtz et al. 2004]) with default settings. Alignment gaps (≥1 kb) and inverted regions (≥10 kb) were extracted from the dnadiff 1coords output file. To construct circular visualizations of this alignment, we used the Circos software (version 0.62.1; [Krzywinski et al. 2009]). For the overview plot, we initially picked the 8 and 11 largest contigs from the DH14 and RACE1 genomes, based on the corresponding N50 values. For each of these contigs we then extracted any further aligning contigs from the other isolate, for which the sum of all aligned regions (with size ≥2 kb and sequence similarity ≥75%) covered at least 10% of both contigs. For the more detailed view of the large-scale rearrangements, we initially selected the contigs with the observed breakpoints and extracted all aligning contigs from the other isolate, for which the sum of all aligning regions (with size ≥ lkb and sequence similarity ≥75%) covered at least 25% of at least one of the contigs. The circular visualizations also depict gene and TE densities along the genome, which were calculated in 10 kb sliding windows (moving by 1 kb at each step) as fraction of bp within each window that is covered by a gene annotation or TE, respectively. For the linear alignment visualizations of the two largest DH14 contigs, we included all aligning regions of at least 1 kb and plotted the corresponding sequence identities from the MUMmer output together with the corresponding gene and TE densities along those contigs. The detailed view of the local inversions observed within the otherwise syntenic alignment between RACE1 tig00005299 and DH14 scaffold 35 was generated with SyMap (version 4.2; [Soderlund et al. 2011]).

### Secretome and core effectorome analysis

The secretomes of all genomes assayed here were identified based on the presence of a signal peptide as detected with SignalP (version 4.1; [Petersen et al. 2011]) and absence of any transmembrane domain in the mature protein as predicted by TMHMM (version 2.0; [Emanuelsson et al. 2007]). Functional domain annotation of the proteomes was performed with InterProScan (Jones, P. et al. 2014).

To define the core *Blumeria-specific* effectorome, we assembled the genomes of the *formae speciales avenae, dicocci, dactylis, lolii, poae, secalis* and *triticale* using the publicly available raw Illumina reads for the isolates AVE, LIB1609, DAC, LOL, POAE, S1459 and T1-20 (Table S13). The assemblies were carried out using ABySS 2.0.2 (Jackman et al. 2017), and the gene space coverage with BUSCO (Table S14). For the *forma specialis tritici* the reference assembly of the isolate 96224 was used. The resulting contig sequences were *de* novo-annotated with one round of MAKER using the same settings as for the DH14 annotation (described previously, also https://github.com/lambros-f/blumeria_2017).

To remove widely conserved, *non-Blumeria* specific proteins, all predicted secreted proteins were used as query in BLASTP searches (version 2.5.0+) against the NCBI non-redundant protein database (nr) with the e-value threshold of 10e-5. Additionally, to derive the presence of core *Blumeria-specific* SPs, an ortholog search was performed using OrthoFinder and the predicted proteomes of the *formae speciales.* To remove potential bias originating from possible conserved secretion signal peptide sequences, the predicted *Bgh* SPs were inserted in the analysis as mature peptides.

To generate a maximum likelihood-based phylogenetic tree for the SPs, all the *Bgh* DH14 mature peptide sequences were aligned with MAFFT v7.310 (‐‐maxiterate 1000 -localpair; [Katoh and Standley 2013]). Afterwards IQ-TREE multicore version 1.6.beta4 (Chernomor et al. 2016) and ModelFinder (Kalyaanamoorthy et al. 2017) were used to select an optimum substitution model and generate the final ML tree. The substitution model used was VT+R8.

To further assess whether *SPs* or *CSEPs* are located in gene sparse regions, BEDTools (Quinlan and Hall 2010) with the functions complement and closest was utilized to calculate the 5’ and 3’ intergenic space lengths for all genes. The resulting tables were introduced in to R in order to generate the corresponding figure using ggplots2. The corresponding R script is deposited in https://github.com/lambros-f/blumeria_2017. As further control we extracted a set of ascomycete core ortholog genes *(COGs)* based on the BUSCO Ascomycota odb9 hidden Markov models (http://busco.ezlab.org/datasets/ascomycota_odb9.tar.gz).

### Divergence landscapes of transposable elements

To generate divergence landscapes for the TEs of the *Letiomycete* fungi, repeat elements were identified in all genomes using RepeatMasker (version 4.0.7, http://www.repeatmasker.org/) with default parameters and *fungi* as the query species. Afterwards the RepeatMasker align output (.aln) was parsed using previously described Perl scripts (https://github.com/4ureliek/Parsing-RepeatMasker-Outputs, [Kapusta et al. 2017]). The selection of genomes used for this analysis (Table S13) and their relation to *Bgh* was derived from the orthology analysis of their proteomes using OrthoFinder (Emms and Kelly 2015).

For the analysis of the dicot infecting powdery mildews the publicly available assemblies were used (Table S13), or in the case of *G. orontii* isolate MGH1 the PacBio reads were assembled with Canu as described previously.

### Duplicate gene search and copy number variation analysis

In order to assess whether duplicate genes exist in the *Bgh* DH14 and RACE1 genomes, MCScanX was used (Wang et al. 2013) with the default parameters. Subsequent analysis to derive copy number variation in all *formae speciales* and their corresponding isolates was carried out as follows. All genomic reads were first quality trimmed using Trimmomatic (version 0.36; [Bolger et al. 2014]) and then aligned to the DH14 genome using BWA-MEM (Li and Durbin 2009). The resulting bam file was sorted using Picard (http://broadinstitute.github.io/picard) and the read depth per bp was extracted using BEDTools (Quinlan and Hall 2010). The copy number of each SP was calculated by the average per bp coverage of the gene model by the respective mapped reads, divided by the average coverage of all 805 SPs using custom R scripts. The distance matrix was computed using the Euclidian method, and the heatmap was generated using heatmap.2 from the package gplots. The bash and R scripts used for this analysis can be found in https://github.com/lambros-f/blumeria_2017.

### Phylogeny of the isolates

The phylogenetic relationship of the *formae speciales* and their corresponding isolates was derived from SNPs. The genomic reads of every isolate were mapped to the DH14 reference genome with BWA-MEM (Li 2013), and the GATK best practices pipeline (DePristo et al. 2011; McKenna et al. 2010) was used for SNP discovery, as previously described (Islam et al. 2016). Afterwards, VCFtools 0.1.15 (Danecek et al. 2011) was deployed with the option –max-missing 1 to keep only common SNPs, resulting in 1,070,264 sites. The resulting VCF files were parsed with custom Perl and bash scripts (https://github.com/lambros-f/blumeria_2017) and imported to SplitsTree (Huson and Bryant 2006) to generate a cladogram based on an UPGMA tree.

### SNP analysis

For isolates A6 and K1, SNPs to DH14 were identified with GATK (DePristo et al. 2011; McKenna et al. 2010) from the BWA-MEM (Li 2013) alignment of short sequence reads as described above. For RACE1, SNPs to DH14 were identified using the nucmer and dnadiff functions of the MUMmer software (version 3.9.4; [Kurtz et al. 2004]) with default settings. Subsequently, for all three isolates, we calculated the SNP frequency as a function of the genomic location by using a 10 kb sliding window that moved 1 kb at each step for all DH14 contigs larger than 50 kb. To further examine the distribution of low and high SNP frequencies, we applied the expectation-maximization (EM) algorithm (function normalmixEM, R-package mixtools) to fit a two-component mixture model to the observed SNP frequencies as described previously (Hacquard et al. 2013; Wicker et al. 2013).

## AUTHOR CONTRIBUTIONS

R.P., P.S.L. and T.M. conceived the study. L.F., B.K., S.K., P.D.S., M.Y.M., S.B. and C.P. performed the experiments. L.F. and B.K. analyzed the data. L.F. and B.K. drafted the manuscript. R.P., P.S.L., P.D.S. and T.M. edited the manuscript.

## ACKNOWLEDGEMENTS

This work was supported by a grant of the Deutsche Forschungsgemeinschaft (DFG)-funded Priority Programme SPP1819 (Rapid evolutionary adaptation - Potential and constraints) to R.P. (PA 861/14-1), the DFG-funded Collaborative Research Centre SFB670/3 (Cell-autonomous Immunity) to P.S.L. (grant #13123509) and the Danish Strategic Research Council, grant no. 10-093504 to C.P. We would like to acknowledge the help of Helder Pedro and of the PhytoPathDB (http://www.phytopathdb.org/) for supporting the re-annotation of the reference isolate DH14. The genome sequencing and assembly data generated in this study have been deposited under the ENA Project IDs PRJEB23502 and PRJEB23162. The new assembly and annotation for the reference Blumeria graminis f.sp. hordei DH14 genome is in addition available through the PhytopathDB database (). The RNA-seq data for DH14 generated in this study has been deposited in the Gene Expression Omnibus (GEO) database, (accession no. GSE106282).

## SUPPLEMENTAL FIGURES

**Supplemental Figure 1:**
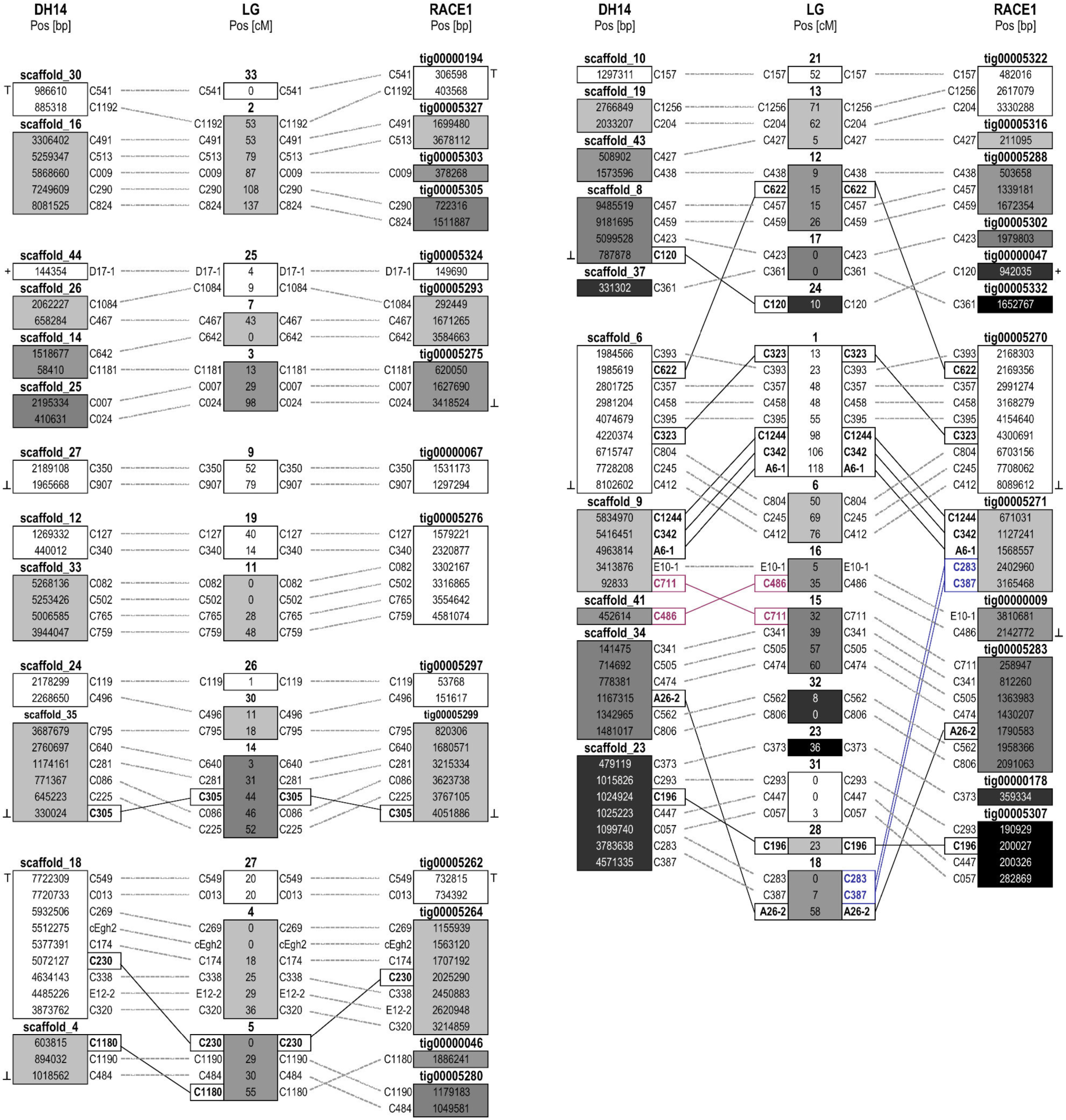
Comparative alignment of the *Bgh* DH14 and RACE1 genome assemblies with a *Bgh* genetic map. The distribution and ordering of 80 single copy EST markers across 30 linkage groups of a previously published genetic map (Pedersen et al. 2002) is visualized in relation to the corresponding genomic locations of these markers in the DH14/RACE1 assemblies. Each box represents a specific genomic contig or linkage group (LG), respectively, and the numbers inside the boxes specify the marker positions on the corresponding contig (in bp) or linkage group (in cM). The corresponding marker identifiers are given next to the boxes. Dashed connector lines represent markers for which the genomic location and genetic map are consistent. Discrepancies between assembly and genetic map are indicated by solid connectors, with black lines representing markers whose location is consistent between assemblies but different from the genetic map, and colored lines representing markers with differences to the genetic map that are specific to either DH14 (dark pink) or RACE1 (blue).

**Supplemental Figure 2:**
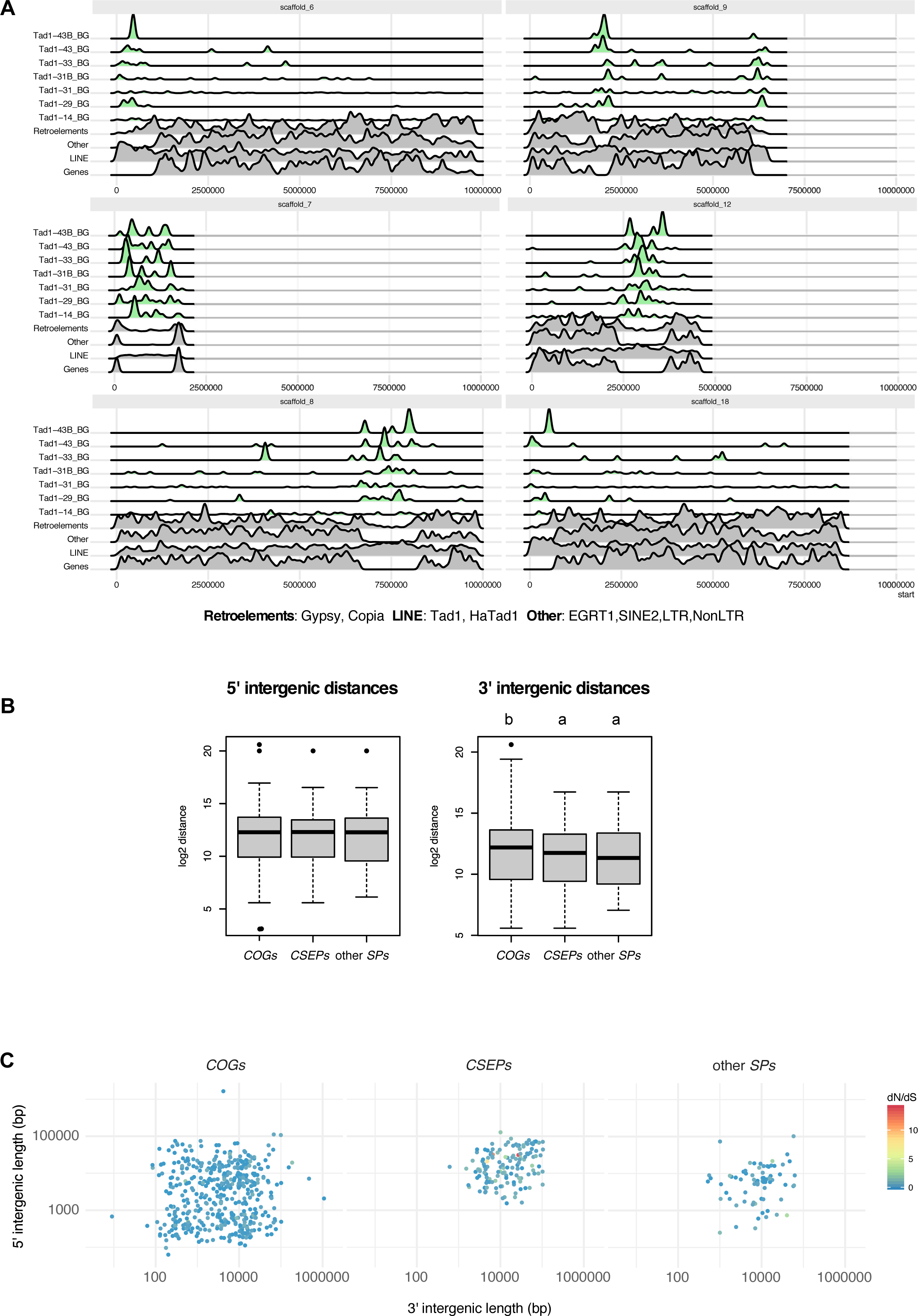
Involvement of TEs in chromosomal organization. (A) Density of different categories of repetitive elements and genes per 50 kb sliding windows in selected scaffolds with putative centromeric regions. A subset of Tad1-like LINE elements that are associated with putative centromeric regions are highlighted in green. (B) Box plots of the 5’ and 3’ intergenic distances for ascomycete core ortholog genes *(COGs), CSEPs* and other secreted protein-coding genes that do not fulfil the *CSEP* criteria ("other SPs"). No statistically significant differences were detected for the 5’ distances (p=0.382; ANOVA) and differing letters indicate statistically significant differences between groups for the 3’ distances (p<0.05; ANOVA with Tukey *post hoc* tests). (C) Plots depicting by color-code the dN/dS ratio of each gene of the three different groups *(COGs, CSEPs,* other *SPs)* in relation to their flanking intergenic length. Genes with dS values of 0 are not shown.

**Supplemental Figure 3:**
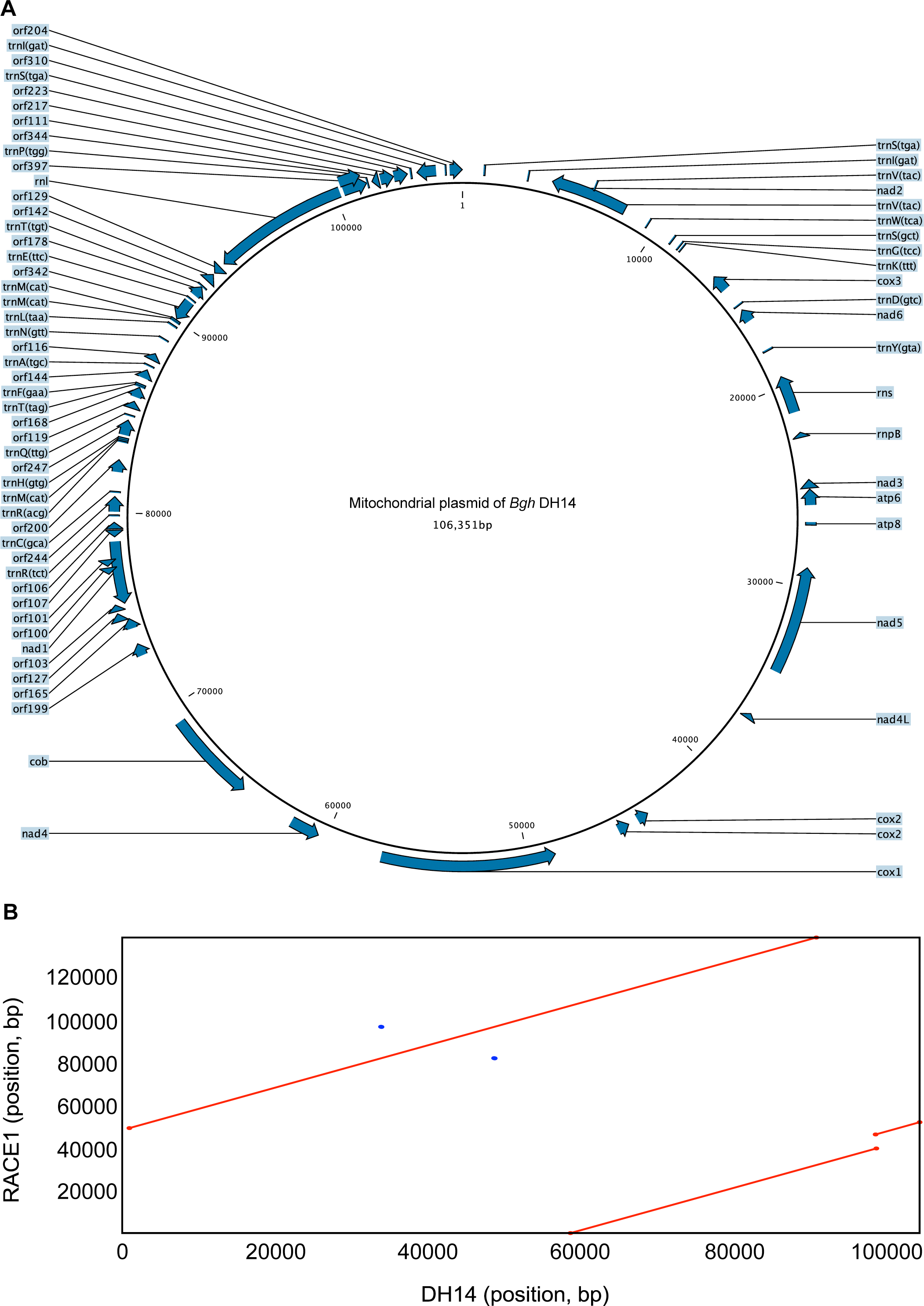
Mitochondrial genomes of *Bgh.* (A) Map and corresponding annotation of the mitochondrial genome of *Bgh* isolate DH14 resulting from an RNAweasel and MFannot run. (B) Nucleotide sequence alignment between the DH14 (x-axis) and RACE1 (y-axis) mtDNA using NUCmer, indicating a putative partial duplication in RACE1.

**Supplemental Figure 4:**
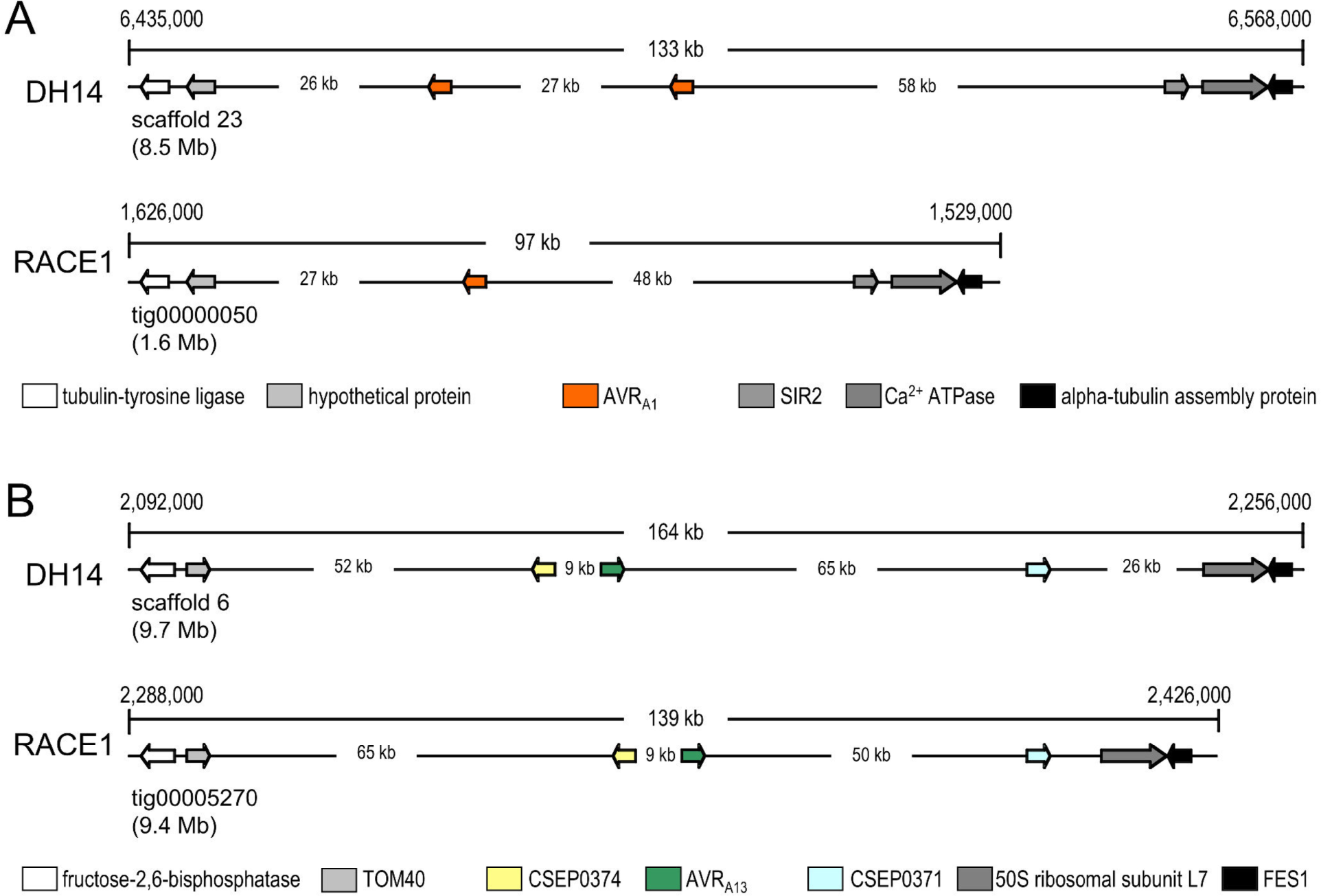
Comparative visualization of the genomic loci harboring *AVR_a1_* and *AVR*_*a13*_ in the *Bgh* isolates DH14 and RACE1. (A) Organization of the genomic locus harboring the previously identified *AVR_a1_* (orange arrows) and some of its flanking genes in DH14 and RACE1. (B) Organization of the genomic locus harboring the previously identified *AVR_a13_* (green arrows) and some of its flanking genes in DH14 and RACE1.

**Supplemental Figure 5:**
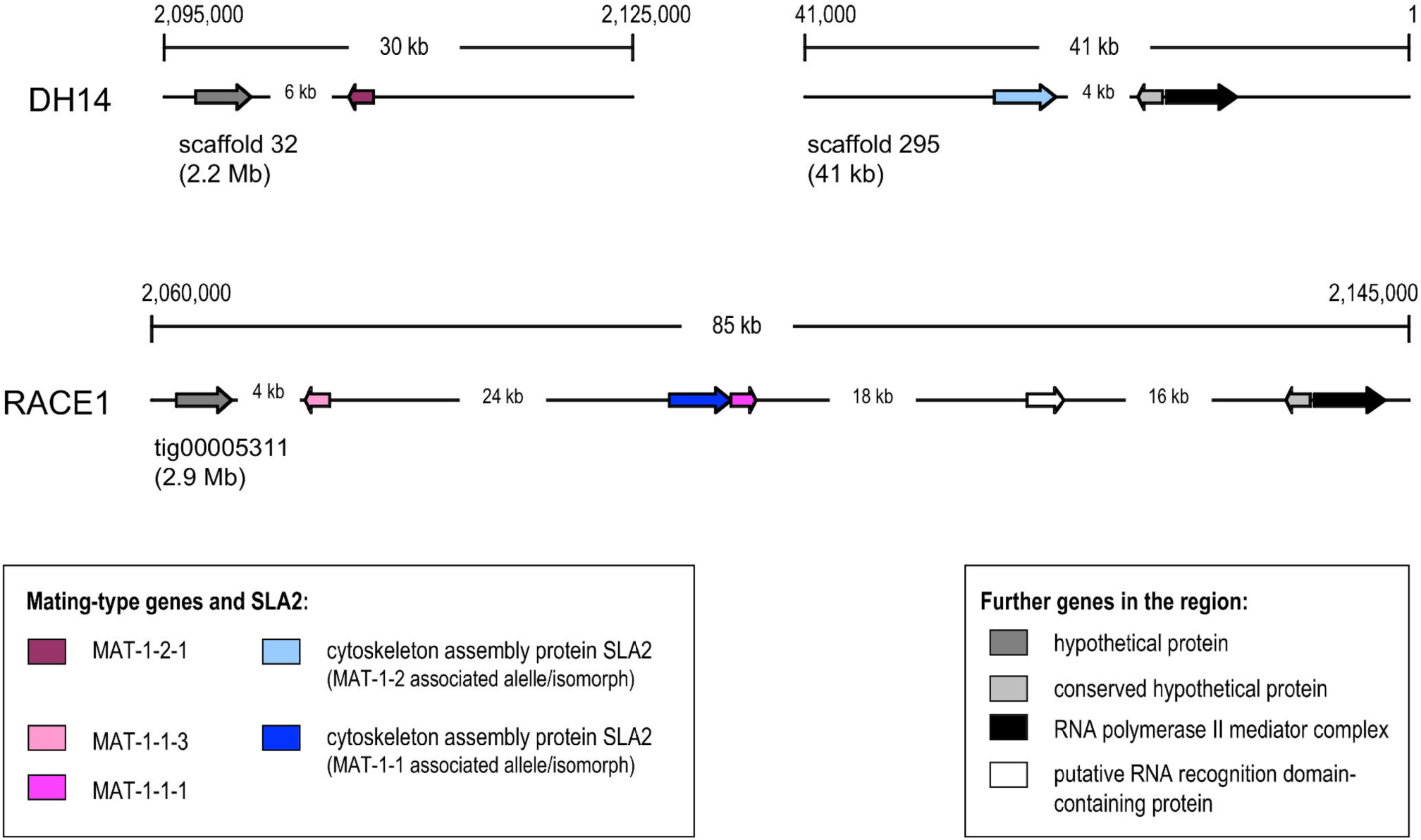
Variation in the mating type locus in the *Bgh* isolates DH14 and RACE1. Organization of the genomic loci containing the mating type genes *(MAT-1-1-1, MAT-1-1-3* and *MAT-1-2-1)* and some of its flanking genes. As DH14 and RACE1 are of opposite mating types, the structure of the mating type locus differs between the two isolates. The genomic locus in RACE1, which is of the MAT-1-1 mating type, was assembled completely, while the respective locus in DH14 (MAT-1-2 mating type) is distributed on two scaffolds.

**Supplemental Figure 6:**
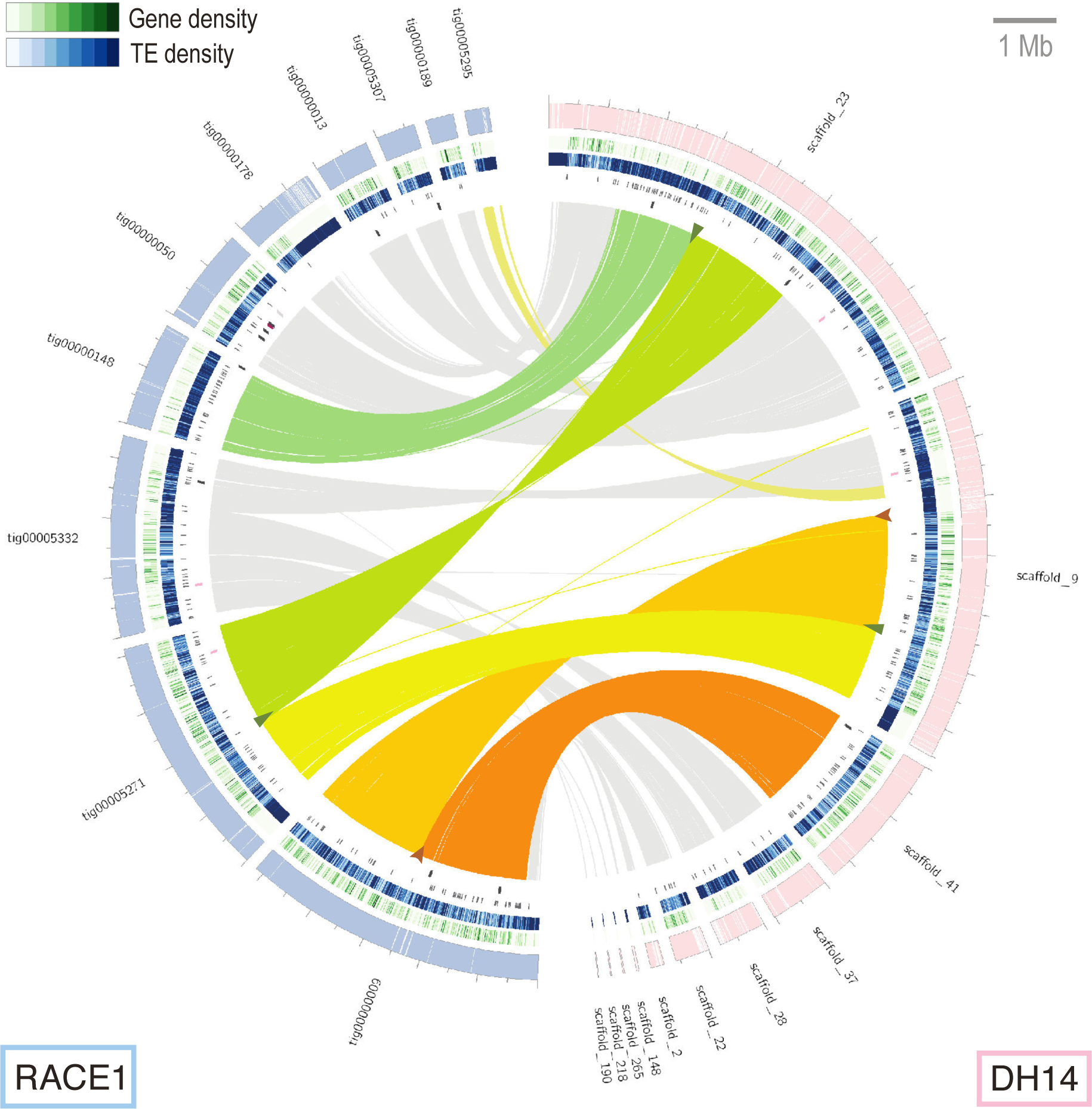
Evidence for two large-scale genomic rearrangements between the isolates DH14 and RACE1. Circos diagram showing evidence for large-scale genomic rearrangements between DH14 and RACE1. The two scaffolds/contigs in the assemblies of DH14 and RACE1 with internal alignment breaks and the corresponding aligning scaffolds/contigs in the other isolate were extracted for visualization. Syntenic regions and alignment breaks were identified based on a whole-genome alignment, and aligning regions of at least 1 kb between the two isolates (with nucleotide sequence similarity ≥75%) are connected with lines in the circular plot. Lines within the syntenic blocks directly flanking the breaks are shown in color while lines in all other blocks are depicted in grey. The positions of the observed alignment breaks are marked by arrowheads colored in green (three breaks likely involved in the same event) and brown (two breaks likely involved in the same event). The surrounding circles represent from the outside: on the right side the DH14 scaffolds (pink) and on the left side the RACE1 (blue), with all unaligned regions (≥ 0.5 kb) indicated as white gaps on the scaffolds/contigs; the gene density (green) and TE density (blue) calculated in 10 kb sliding windows; the locations of all genes predicted to code for SPs; the locations of isolate-specific genes coding for SPs (dark red) or any other proteins (black); and isolate-specific additional gene copies/paralogs coding for SPs (pink) or any other proteins (grey).

**Supplemental Figure 7:**
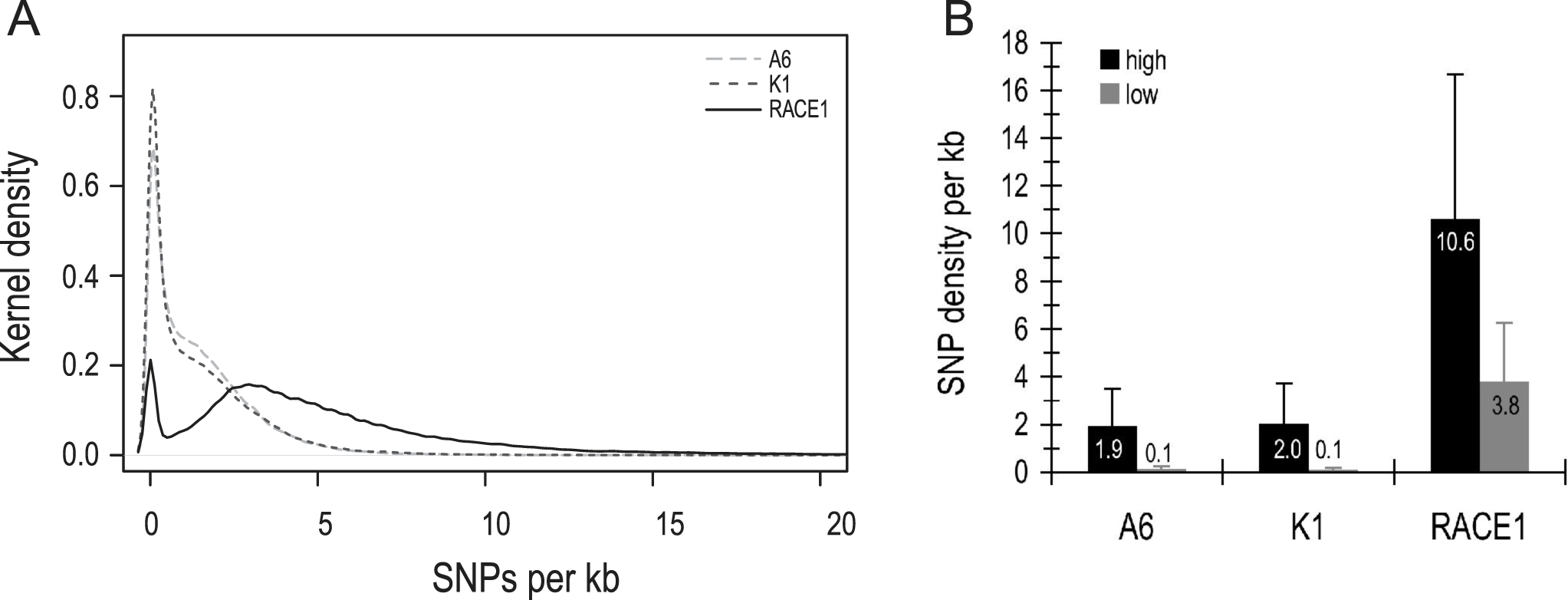
Frequency of single-nucleotide polymorphisms (SNPs) between *Bgh* isolates. (A) Kernel density plot of the SNP frequencies per kb in 10 kb sliding windows, observed for the three *Bgh* isolates A6, K1 and RACE1 relative to the reference isolate DH14. The plot depicts Gaussian kernel density estimates calculated at a smoothing bandwidth of 0.12. (B) Average SNP frequencies for A6, K1 and RACE1 in 10 kb sliding windows of low and high SNP density as estimated by a two-component mixture model that was fitted to the observed SNP frequencies using the expectation-maximization algorithm. Error bars indicate the corresponding standard deviations estimated by the mixture model.

**Supplemental Figure 8:**
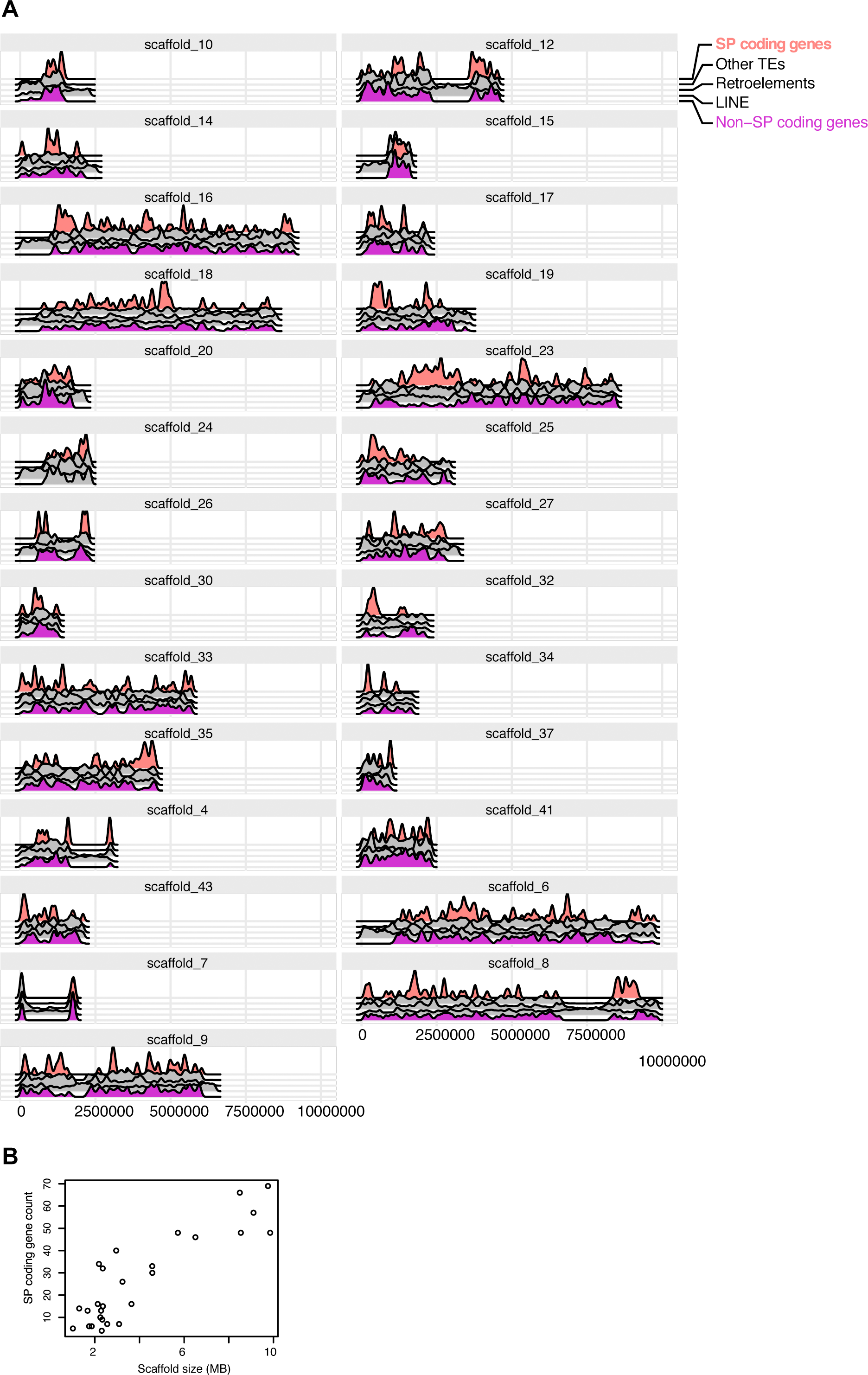
Distribution of SP and non-SP coding genes in *Bgh* DH14 scaffolds larger than 1 MB. (A) Density plots of SP coding genes (orange), non-SP coding genes (purple) and different types of TE elements (gray) in 50 kb sliding windows. Scaffolds depicted here were selected based on their size (>1 MB) and represent ~87% of the total genomic sequence. (B) Number of SP coding genes per scaffold plotted against the respective total scaffold size, showing positive correlation (r=0.88, p<0.001).

**Supplemental Figure 9:**
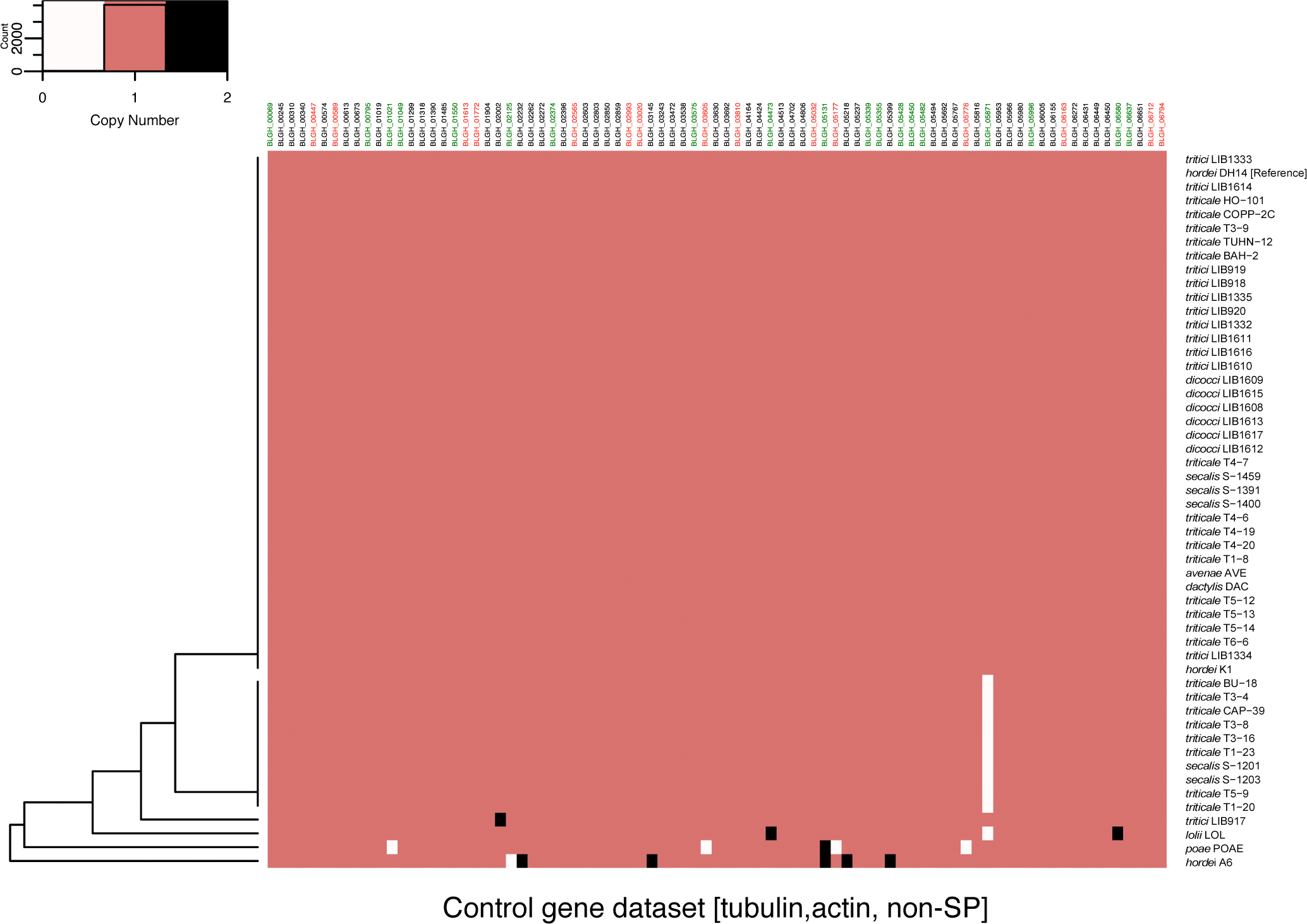
CNV of widely conserved genes between *B. graminis formae speciales.* Heatmap illustrating the copy number of genes with putatively widely conserved functions. Using the same pipeline as for the generation of Figure 3A, all 34 genes with a PFAM annotation including the terms "tubulin" (highlighted in red) or "actin" ((highlighted in green) and 49 genes coding for non-SP genes with conserved domains were used as a control dataset to estimate the error rate of the CNV calling pipeline. The heatmap depicts the color-coded copy number of these genes per individual genome of various *B. graminis formae speciales (avenae, dactylis, dicocci, hordei, lolii, poae, secalis, triticale* and *tritici*), each represented by one or more isolates as indicated on the right. The dendrogram on the left is based on the hierarchical clustering (Euclidean method) of the CNV values for every dataset.

**Supplemental Figure 10:**
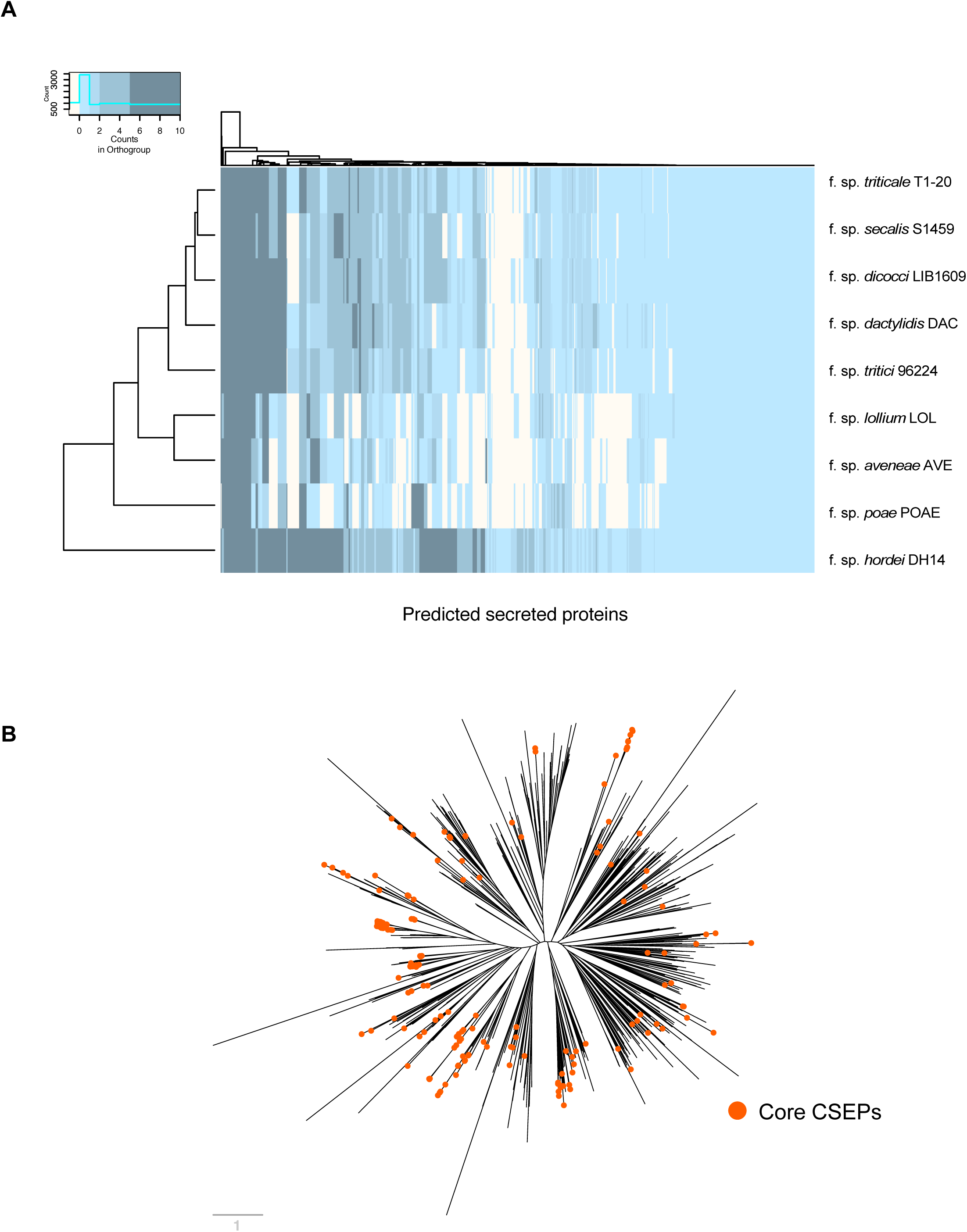
Secretome orthology relations and core effectorome phylogeny. (A) Heatmap of *SP* orthologs found for the *formae speciales* genomes after ortholog clustering using OrthoFinder on the predicted proteomes of the isolates T1-20, S1459, LIB1609, DAC, 96224, LOL, AVE, POAE, DH14. Every column corresponds to one of the 805 *Bgh* DH14 predicted *SPs,* while color-coding depicts the number of orthologs in the corresponding orthogroup. Hierarchical clustering (Euclidean method) for the *formae speciales* and the *SPs* are given on the left and the top of the heatmap, respectively. (B) Maximum likelihood phylogeny tree of the 805 SPs. The tree was generated using IQ-TREE based on the mature peptide sequences of the *Bgh* DH14 SPs. Orange edge tips indicate the 190 core CSEPs which have orthologs in all *formae speciales.* The scale bar indicates the number of amino-acid substitutions per site.

**Supplemental Figure 11:**
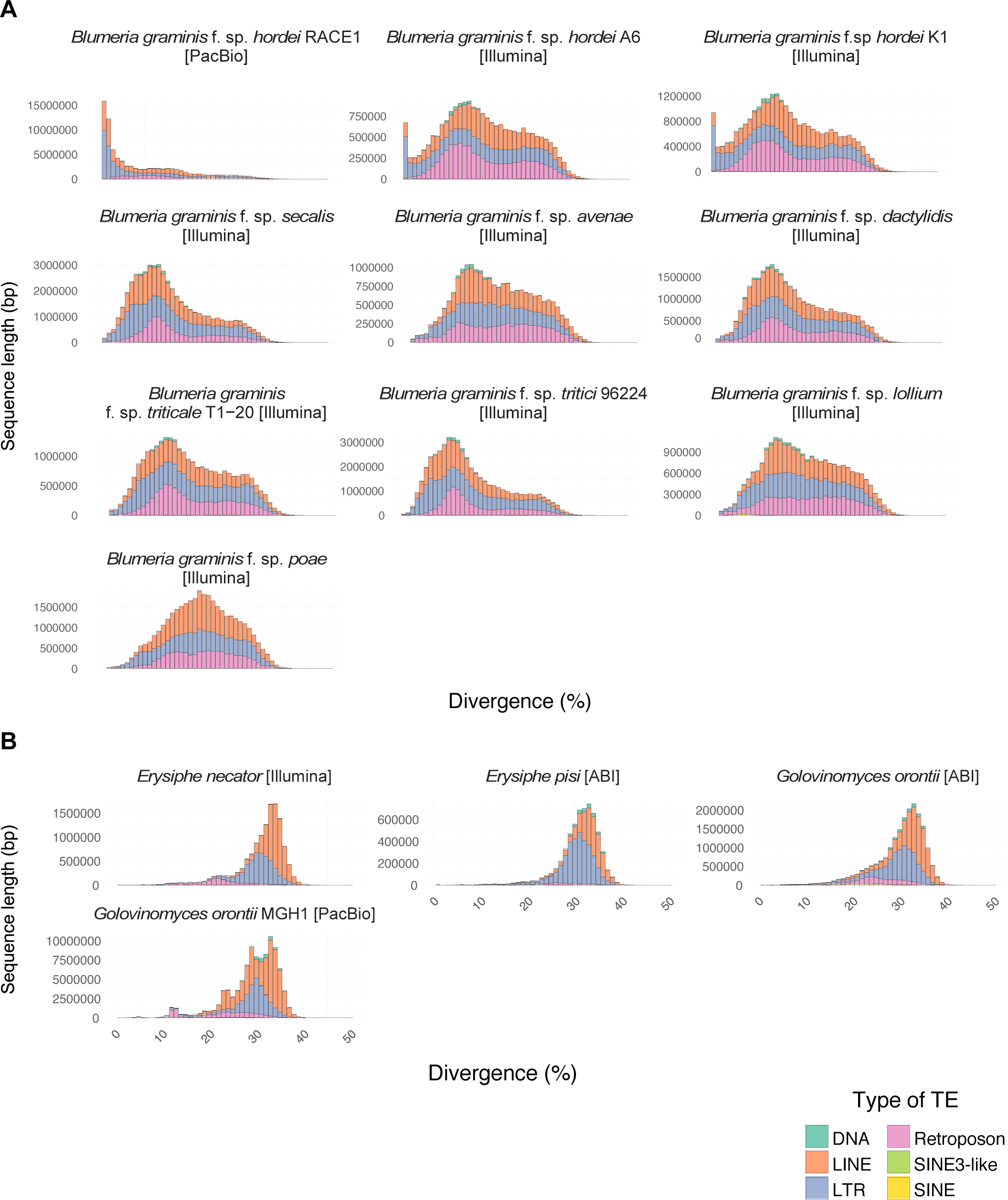
Representatives of the genus *Blumeria* show less TE divergence than representatives of the genera *Erishyphe* and *Golovinomyces.* (A) The histograms indicate the frequency of a given sequence divergence for TE families of 10 *B. graminis* genomes. The genomes, which were assembled based on various sequencing platforms (PacBio or Illumina), were surveyed for their repeat content and repeat landscapes for each genome based on % nucleotide divergence to the consensus TE sequences were calculated out of the RepeatMasker output using Perl scripts. Sequence divergence (x-axis) is plotted against frequency (number of sequences; y-axis) for each of the genomes. (B) The histograms indicate the frequency of a given sequence divergence for TE families of 3 dicot-infecting powdery mildew species (*Erysiphe pisi, E. necator* and *Golovinomyces orontii*). The genomes, which were assembled based on various sequencing platforms (PacBio, ABI Solid or Illumina), were surveyed for their repeat content and repeat landscapes for each genome based on % nucleotide divergence to the consensus TE sequences were calculated out of the RepeatMasker output using Perl scripts. Sequence divergence (x-axis) is plotted against frequency (number of sequences; y-axis) for each of the genomes.

## Abbreviations

*AVR*: avirulence effector gene
*Bgh*: *Blumeria graminis* f.sp. *hordei*
CNV: copy number variation
CSEP: candidate secreted effector protein
EST: expressed sequence tag
FPKM: fragments per kilobase [sequence length] and million [sequenced fragments]
mtDNA: mitochondrial DNA
SNP: single nucleotide polymorphism
SP: secreted protein
TE: transposable element

